# LEAP2 is a more conserved ligand than ghrelin for fish GHSRs

**DOI:** 10.1101/2022.09.21.508860

**Authors:** Hao-Zheng Li, Xiao-Xia Shao, Ya-Fen Wang, Ya-Li Liu, Zeng-Guang Xu, Zhan-Yun Guo

## Abstract

Recently, liver-expressed antimicrobial peptide 2 (LEAP2) was identified as an endogenous competitive antagonist and an inverse agonist of the ghrelin receptor GHSR. However, its functions in lower vertebrates are not well understood. Our recent study demonstrated that both LEAP2 and ghrelin are functional towards a fish GHSR from *Latimeria chalumnae*, an extant coelacanth believed to be one of the closest ancestors of tetrapods. However, amino acid sequence alignment identified that the 6.58 position (Ballesteros-Weinstein numbering system) of most fish GHSRs are not occupied by an aromatic Phe residue, which is absolutely conserved in all known GHSRs from amphibians to mammals, and is responsible for human GHSR binding to its agonist, ghrelin. To test whether these unusual fish receptors are functional, we studied the ligand binding properties of three representative fish GHSRs, two from *Danio rerio* (zebrafish) and one from *Larimichthys crocea* (large yellow croaker). After overexpression in human embryonic kidney 293T cells, the three fish GHSRs retained normal binding to all tested LEAP2s, except for a second LEAP2 from *L. crocea*. However, they displayed almost no binding to all chemically synthesized n-octanoylated ghrelins, despite these ghrelins all retaining normal function towards human and coelacanth GHSRs. Thus, it seems that LEAP2 is a more conserved ligand than ghrelin towards fish GHSRs. Our results not only provided new insights into the interaction mechanism of GHSRs with LEAP2s and ghrelins, but also shed new light on the functions of LEAP2 and ghrelin in different fish species.

## 1. Introduction

Liver-expressed antimicrobial peptide 2 (LEAP2 or LEAP-2) was first identified in 2003 and was named according to its weak antimicrobial activity measured using *in vitro* assays [1]. Mature LEAP2 is derived from a prepropeptide that is ubiquitously present in all vertebrates and is highly conserved in evolution. Only recently was the main biological function of LEAP2 disclosed: It is actually an endogenous competitive antagonist and an inverse agonist of the growth hormone secretagogue receptor (GHSR) [2−6]. As a formerly orphan A-class G protein-coupled receptor (GPCR), GHSR was first identified as the receptor of certain synthetic growth hormone secretagogues in 1996, hence its name [7]. Soon after, the gastric peptide ghrelin was identified as its endogenous agonist, thus GHSR is also known as the ghrelin receptor [8]. In some species, two transcripts are produced from *GHSR* gene: One variant encodes a functional longer isoform 1a (GHSR1a) with seven transmembrane domains (TMDs), while the other variant encodes a non-functional shorter isoform 1b (GHSR1b) with five TMDs. In the present study, all GHSRs refer to the functional longer isoform. Mature ghrelin is derived from precursors encoded by *GHRL* gene, and carries an essential *O*-fatty acyl moiety, typically n-octanoyl, at the side-chain of its third Ser residue [8]. This special posttranslational modification is catalyzed by membrane bound *O*-acyltransferase domain containing 4 (MBOAT4), also known as ghrelin *O*-acyltransferase (GOAT) [9,10]. The *LEAP2, GHSR, GHRL*, and *MBOAT4* genes are ubiquitously present in all vertebrates from fish to mammals, suggesting that the ghrelin-LEAP2-GHSR system probably originated in ancient fish and then spread to all vertebrate lineages during evolution.

In mammals, LEAP2 and ghrelin exert important functions in energy metabolism and cellular homeostasis mediated by their receptor, GHSR [11−15]. However, their functions in lower vertebrates are not well understood. Our recent study demonstrated that both LEAP2 and ghrelin are functional towards a fish GHSR from *Latimeria chalumnae*, a ‘living fossil’ coelacanth believed to be one of the closest ancestors of tetrapods [16]. In all known GHSRs from amphibians to mammals, their 6.58 position (according to the Ballesteros-Weinstein generic numbering system) at the extracellular end of TMD6 is always occupied by an aromatic Phe residue [17,18]. However, only a few fish GHSRs, such as that from the extant coelacanth *L. chalumnae*, have a Phe residue at their 6.58 position (Fig. S1). For some other fish GHSRs, such as the two receptors from *Danio rerio* (zebrafish), their 6.58 position is occupied by a large aliphatic residue, such as Ile, Val, or Met (Fig. S1). For most other fish GHSRs, such as that from *Larimichthys crocea* (large yellow croaker), their 6.58 position is occupied by a hydrophilic Gln residue, or occasionally by a hydrophilic His residue (Fig. S1).

In human GHSR, the highly conserved Phe6.58 (Phe286) is essential for receptor binding to ghrelin, but not for receptor binding to LEAP2 [18,19]. Thus, we were curious as to whether those fish GHSRs without a Phe residue at their 6.58 position are functional. To answer this question, in the present study, we examined the ligand-binding properties of three representative fish GHSRs, two from *D. rerio* and one from *L. crocea*. Our results demonstrated that the three fish GHSRs retained normal binding to all tested LEAP2s, except to a second LEAP2 from *L. crocea*, but they displayed almost no binding to all synthetic n-octanoylated ghrelins tested, despite these ghrelins being fully functional towards human and coelacanth GHSRs. Thus, both *D. rerio* and *L. crocea* have a functional LEAP2 displaying high activity, not only towards their cognate GHSRs, but also towards those receptors from human and coelacanth. However, it seems that ghrelins cannot bind to and activate the fish GHSRs from *D. rerio* and *L. crocea*, but are fully active towards GHSRs from humans and coelacanth.

## 2. Experimental procedures

### 2.1. Recombinant expression of LEAP2s

The expression constructs for N-terminally 6×His-tagged precursors of human LEAP2 (designated as Hs-LEAP2) and *L. chalumnae* LEAP2 (Lc-LEAP2) were generated in our previous studies [3,18]. For the mature LEAP2 from *D. rerio* (Dr-LEAP2) and the mature LEAP2s from *L. crocea* (La-LEAP2A and La-LEAP2B), their coding DNA fragments were chemically synthesized at GeneWiz (Suzhou, China) using *Escherichia coli*-biased codons, and ligated into a pET vector, resulting in the expression constructs encoding N-terminally 6×His-tagged precursors (Fig. S2). Alternatively, the coding region of Dr-LEAP2 and La-LEAP2A were also fused to the downstream of human C4ORF48 coding sequence, and then inserted into the pET vector, resulting in the expression constructs encoding N-terminally 6×His-tagged and C4ORF48-fused precursors (Fig. S2). A sortase A recognition motif was introduced to the C-terminus of 6×His-Hs-LEAP2 and 6×His-C4ORF48-La-LEAP2A for preparation of the NanoLuc-based binding tracers (Fig. S2).

All LEAP2 precursors were overexpressed in the *E. coli* strain BL21(DE3) and their expression was determined by sodium dodecyl sulfate-polyacrylamide gel electrophoresis (SDS-PAGE). Thereafter, the overexpressed precursors in the inclusion bodies were solubilized via an *S*-sulfonation approach, and purified by an immobilized metal ion affinity chromatography (Ni^2+^ column). For the 6×His-tagged precursors, the eluted fraction from the Ni^2+^ column was directly subjected to in vitro refolding according to our previous procedure for refolding of 6×His-Hs-LEAP2 [3,18]. After purification via high performance liquid chromatography (HPLC) using a semi-preparative C_18_ reverse-phase column (Zorbax 300SB-C18, 9.4 × 250 mm; Agilent Technologies, Santa Clara, CA, USA), the refolded 6×His-tagged precursors were treated with enterokinase (New England Biolabs, Ipswich, MA, USA) to remove their N-terminal 6×His-tag according to our previous procedure [3,18] and purified by HPLC using an analytical C_18_ reverse-phase column (Zorbax 300SB-C18, 4.6 × 250 mm; Agilent Technologies). For those C4ORF48-fused LEAP2 precursors, the eluted fraction from the Ni^2+^ column was dialyzed against water overnight, and the precipitate was collected and dissolved in endoproteinase Lys-C digestion buffer (100 mM Tris-Cl, pH8.5, 3 M urea) at the final concentration of ∼5 mg/ml. After addition of Lys-C endoproteinase (Yaxin Bio, Shanghai, China) to the final concentration of ∼5 μg/ml, the digestion reaction was conducted at 30 °C overnight. Thereafter, the digestion mixture was loaded onto a spin Ni^2+^ column and the flow-through fraction was subjected to in vitro refolding according to the procedure for refolding 6×His-Hs-LEAP2 [3,18]. The refolded mature LEAP2s were then purified by HPLC using an analytical C_18_ reverse-phase column (Zorbax 300SB-C18, 4.6 × 250 mm; Agilent Technologies). Purity of the mature LEAP2s was analyzed by HPLC and their identity was confirmed by electrospray mass spectrometry.

### 2.2. Chemical synthesis of ghrelins and UAGs

The mature n-octanoylated human ghrelin (Hs-ghrelin), *L. chalumnae* ghrelin (Lc-ghrelin), *D. rerio* ghrelins (Dr-ghrelin-22 and Dr-ghrelin-19), and *L. crocea* ghrelin (La-ghrelin-23), as well as some unacylated forms (Dr-UAG-22 and La-UAG-23) were chemically synthesized at GL Biotech (Shanghai, China) via solid-phase peptide synthesis using standard Fmoc methodology. The crude synthetic peptides were purified to homogeneity by HPLC sequentially using a semi-preparative C_18_ reverse-phase column and an analytical C_18_ reverse-phase column (Zorbax 300SB-C18, 9.4 or 4.6 × 250 mm; Agilent Technologies). Purity of the synthetic peptides was analyzed by HPLC and their identity was confirmed by electrospray mass spectrometry.

### 2.3. Generation of expression constructs for GHSRs

The expression constructs for human GHSR (Hs-GHSR) and *L. chalumnae* GHSR (Lc-GHSR) were generated in our previous studies [3,16]. The coding regions of *D. rerio* GHSRA and GHSRB (Dr-GHSRA and Dr-GHSRB) as well as *L. crocea* GHSR (La-GHSR) were chemically synthesized at TsingKe Biological Technology (Beijing, China) according to their published nucleotide sequences at National Center for Biotechnology Information (NCBI). After cleaved by restriction enzymes KpnI and AgeI, the synthetic DNA fragments were cloned into pcDNA6 vectors either carrying the coding sequence of a secretory large NanoLuc fragment (sLgBiT) or not. The resultant mammalian cell expression constructs either encode an N-terminally sLgBiT-fused GHSR, or a GHSR without any tags. The coding region of these fish GHSRs was confirmed by DNA sequencing.

### 2.4. Preparation of the SmBiT-based tracers and the NanoLuc-based tracers

The SmBiT-based binding tracers, ghrelin-SmBiT and LEAP2-SmBiT, were generated in our previous studies via chemical conjugation of a C-terminally Cys-tagged synthetic SmBiT with a C-terminally Cys-tagged synthetic Hs-ghrelin or a recombinant Hs-LEAP2 mutant [3,18].

The NanoLuc-based binding tracers, Hs-LEAP2-Luc and La-LEAP2A-Luc, were prepared via sortase A-catalyzed ligation of the C-terminally sortase A recognition motif-fused LEAP2s (Hs-LEAP2-srt and La-LEAP2-srt) with an N-terminally 6×Gly-fused NanoLuc according to our previous procedures for ligation of Nanoluc with other peptides [20]. The circular sortase A was overexpressed in *E. coli* via an intein-mediated cyclization and purified according to our previous procedure [21]. The coding DNA sequence of the 6×Gly-fused NanoLuc was amplified by polymerase chain reaction (PCR) and then ligated into a pET vector, resulting in an expression construct encoding a NanoLuc carrying tandem 6×His-tag and 6×Gly at its N-terminus (Fig. S3). After overexpressed in *E. coli*, soluble 6×His-6×Gly-NanoLuc protein was purified by a Ni^2+^ column, and the eluted fraction was dialyzed against enterokinase digestion buffer (50 mM Tris-Cl, pH8.5, 2.0 mM CaCl_2_, 50 mM NaCl). Thereafter, enterokinase stock (New England Biolabs) were added (∼1 μl of enterokinase stock versus ∼5 mg of the NanoLuc precursor). After incubation at 25 °C overnight, the reaction mixture was applied to a DEAE ion-exchange column (TSKgel DEAE-5PW, 7.5 mm × 75 mm; Sigma-Aldrich, St. Louis, MO, USA), and mature 6×Gly-NanoLuc was eluted from the column by a linear NaCl gradient and confirmed by SDS-PAGE and bioluminescence measurement. To conduct ligation, the purified 6×Gly-NanoLuc, circular sortase A, and the ligation version LEAP2 were mixed together at the final concentrations of ∼25 μM each in the ligation buffer (100 mM Tris-Cl, pH8.0, 5.0 mM CaCl_2_, 1.0 M urea). After overnight incubation at 30 °C, the ligation mixture was applied to a DEAE ion-exchange column (TSKgel DEAE-5PW, 7.5 mm × 75 mm; Sigma-Aldrich), and the ligation product was eluted from the column by a linear NaCl gradient, confirmed by SDS-PAGE and bioluminescence measurement.

### 2.5. The NanoBiT-based homogenous binding assays

The NanoBiT-based homogenous binding assays were conducted according to our previous procedures [3,16–18]. The expression constructs for the sLgBiT-fused GHSRs were transiently transfected into human embryonic kidney (HEK) 293T cells, respectively. The next day, the transfected cells were trypsinized, seeded into white opaque 96-well plates, and continuously cultured for ∼24 h to ∼90% confluence. To conduct binding assays, the medium was removed and binding solution [phosphate-buffered saline (PBS) plus 0.1% bovine serum albumin (BSA) and 0.01% Tween-20] was added (50 μl/well). For saturation binding assays, the binding solution contained varied concentrations of ghrelin-SmBiT or LEAP2-SmBiT. For competition binding assays, the binding solution contained a constant concentration of LEAP2-SmBiT and varied concentrations of competitor. After incubation at 24 °C for ∼1 h, 20-fold diluted NanoLuc substrate (Promega, Madison, WI, USA) was added (diluted by PBS, 10 μl/well) and bioluminescence was immediately measured on a SpectraMax iD3 plate reader (Molecular Devices, Sunnyvale, CA, USA). The measured bioluminescence data were expressed as mean ± standard deviation (SD, *n* = 3) and fitted to one-site binding model using SigmaPlot 10.0 software (SYSTAT software, Chicago, IL, USA).

### 2.6. The washing-based binding assays

The washing-based binding assays were conducted according to our previous procedures [3,20]. The expression constructs for the untagged GHSRs were transiently transfected into HEK293T cells, respectively. The next day, the transfected cells were trypsinized, seeded into white opaque 96-well plates, and continuously cultured for ∼24 h to ∼90% confluence. To conduct the binding assays, the medium was removed and binding solution (PBS plus 1% BSA) was added (50 μl/well). For competition binding assays, the binding solution contained a constant concentration of the NanoLuc-ligated LEAP2 tracer and varied concentrations of competitor. After incubation at 24 °C for ∼1 h, the binding solution was removed and the cells were washed twice with ice-cold PBS (200 μl/well for each washing). Thereafter, 100-fold diluted NanoLuc substrate was added (diluted by PBS, 50 μl/well), and bioluminescence was immediately measured on a SpectraMax iD3 plate reader (Molecular Devices). The measured bioluminescence data were expressed as mean ± SD (*n* = 3) and fitted to sigmoidal or linear curves using SigmaPlot 10.0 software (SYSTAT software).

### 2.7. Receptor activation assays

The GHSR activation assays were conducted according to our previous procedures [3,16–18]. The expression constructs for the untagged GHSRs were transiently transfected into HEK293T cells together with a cAMP response element (CRE)-controlled NanoLuc reporter vector pNL1.2/CRE. The next day, the transfected cells were trypsinized, seeded into white opaque 96-well plates, and continuously cultured for ∼24 h to ∼90% confluence. To conduct the activation assays, the medium was removed and activation solution (serum-free DMEM medium plus 1% BSA) was added (50 μl/well). The activation solution contained varied concentrations of ghrelins or UAGs. After the cells were continuously cultured at 37 °C for ∼4 h, 20-fold diluted NanoLuc substrate was added (diluted by PBS, 10 μl/well) and bioluminescence was immediately measured on a SpectraMax iD3 plate reader (Molecular Devices). The measured bioluminescence data were expressed as mean ± SD (*n* = 3) and fitted to sigmoidal or linear curves using SigmaPlot 10.0 software (SYSTAT software).

## Results

### 3.1. GHSRs, LEAP2s, and ghrelins from D. rerio and L. crocea

According to the published sequence data at the NCBI, *D. rerio* has two *ghsr* genes, *ghsra* (NM_001146272) and *ghsrb* (XM_002666671), which encode two receptors (Dr-GHSRA and Dr-GHSRB) with typical seven TMDs; however, *L. crocea* has only one *ghsr* gene (XM_002666671) that also encodes an A-class GPCR (La-GHSR). As shown in Fig. 1A, the three fish GHSRs share relatively high overall sequence similarity with human GHSR (Hs-GHSR) and the coelacanth GHSR from *L. chal<u>u</u>mnae* (Lc-GHSR); however, their 6.58 position is not occupied by an aromatic Phe residue, which is absolutely conserved in all known GHSRs from amphibians to mammals. To test whether these unusual fish GHSRs retain a normal response to ghrelin and LEAP2, we chemically synthesized their full-length coding regions and generated mammalian cell expression constructs that encode these fish GHSRs with or without an N-terminal sLgBiT. The sLgBiT-fused receptors were used for NanoBiT-based ligand-binding assays, while the untagged receptors were used for activation assays and washing-based ligand-binding assays.

**Fig. 1.**
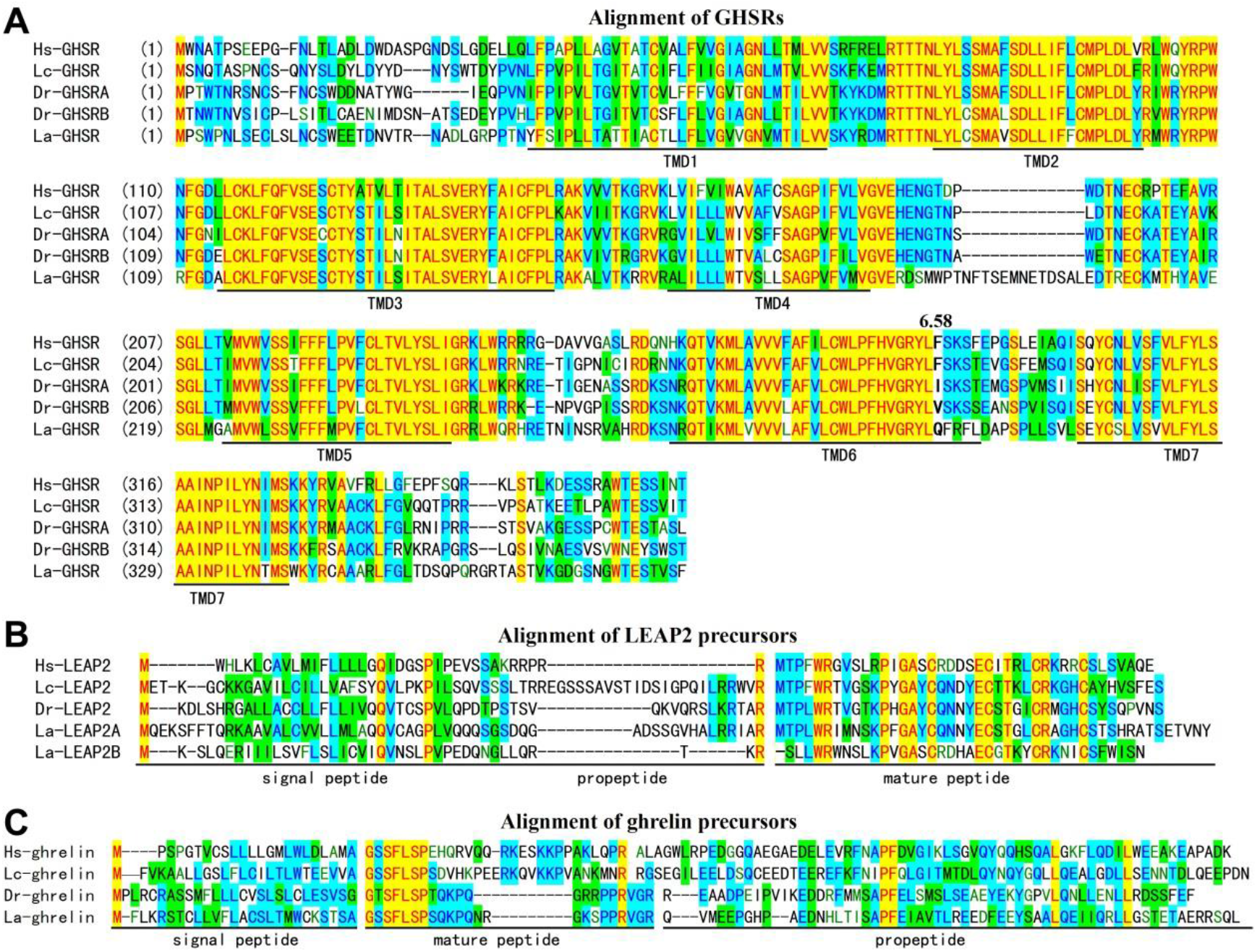
Amino acid sequence alignment of GHSRs (**A**), LEAP2 precursors (**B**), and ghrelin precursors (**C**) from *H. sapiens, L. chalumnae, D. rerio*, and *L. crocea*. The TMDs of GHSRs are underlined and labeled, the residues at 6.58 position are shown as bold letters and indicated. The signal peptide, propeptide, and mature peptide of LEAP2 precursors and ghrelin precursors are indicated.

*D. rerio* has one *leap2* gene (NM_001128777), whereas *L. crocea* has two *leap2* genes, *leap2a* (NM_001303342) and *leap2b* (XM_010736581). These *leap2* genes both encode LEAP2 precursors containing an N-terminal signal peptide, a short propeptide, and a mature peptide (Fig. 1B). After post-translational processing, mature LEAP2s with two disulfide bonds are expected to be released (Fig. 1B). Some N-terminal residues, such as Phe4, Trp5, and Arg6, are important for the binding of human LEAP2 (Hs-LEAP2) to its receptor [18]. These important residues are all present in Lc-LEAP2 (Fig. 1B), whereas the aromatic Phe4 is replaced by a large aliphatic Leu residue in the LEAP2s from *D. rerio* and *L. crocea* (Fig. 1B). Moreover, La-LEAP2B lacks the N-terminal Met residue compared with other LEAP2s (Fig. 1B). It is unknown whether these fish LEAP2s are functional towards their cognate receptors as well as other GHSRs. In the present study, we prepared these fish LEAP2s by overexpression of a suitable precursor in *E. coli* and subsequent *in vitro* processing. The resultant mature LEAP2s all displayed the expected molecular mass, as measured using mass spectrometry (Table 1) and showed a single elution peak on HPLC (Fig. S4A).

**Table 1.**
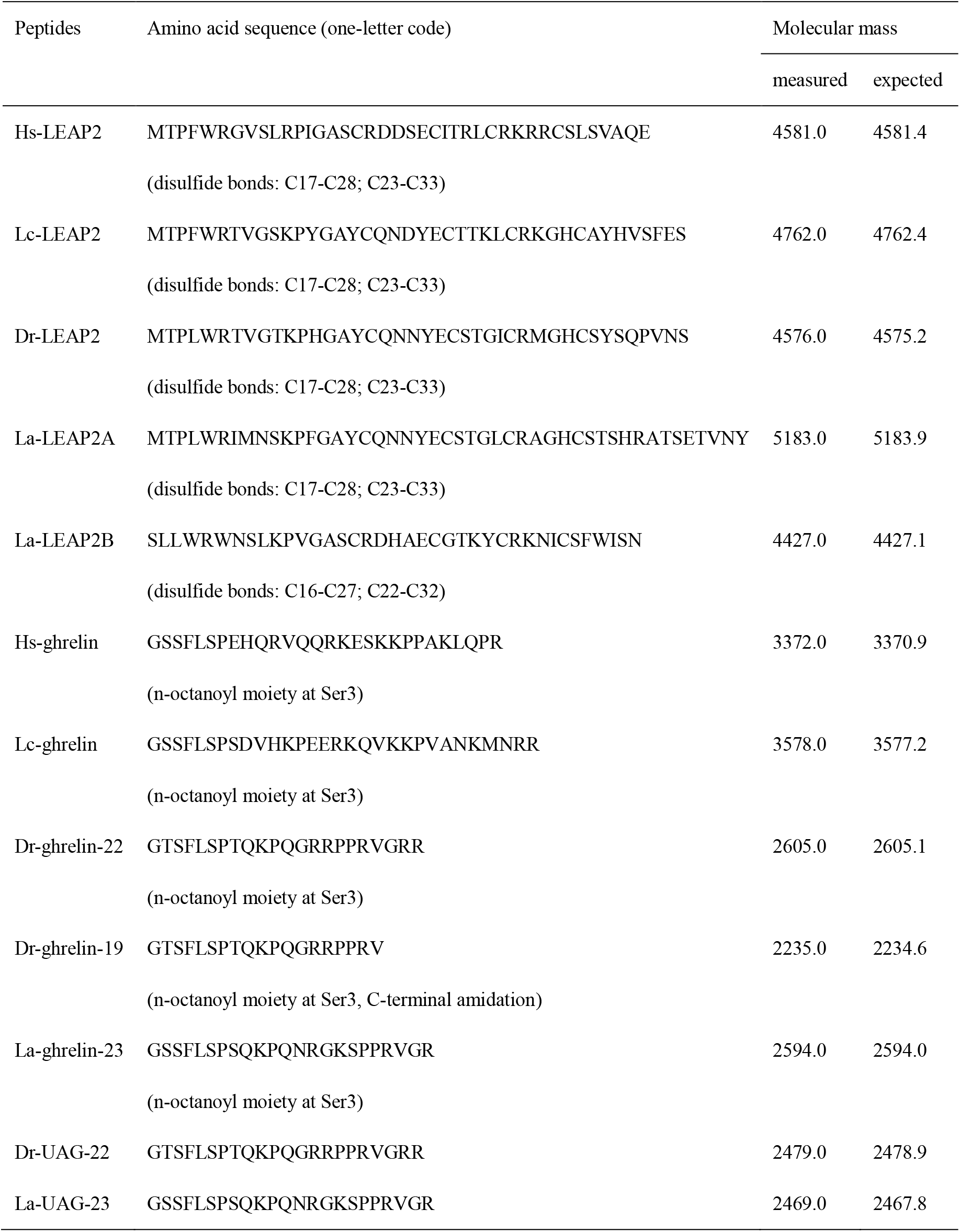
Summary of the recombinant LEAP2s and synthetic ghrelins used in the present study.

Both *D. rerio* and *L. crocea* have a single *ghrl* gene (XM_021476948 and NM_001303331, respectively) that encodes a ghrelin precursor containing an N-terminal signal peptide, a mature peptide, and a C-terminal propeptide (Fig. 1C). The expected mature peptides from *D. rerio* (Dr-ghrelin) and *L. crocea* (La-ghrelin) are shorter than the orthologs from *H. sapiens* and *L. chalumnae*; however, their six N-terminal residues are highly conserved, including a Ser residue at the third position for the expected *O*-fatty acyl modification. Both *D. rerio* and *L. crocea* have an *mboat4* gene (NM_001122944 and XM_027273217, respectively) sharing high sequence similarity with the orthologs from human and coelacanth (Fig. S5). To test the activity of these fish ghrelins towards their cognate receptors and other GHSRs, we chemically synthesized the n-octanoylated fish ghrelins and some unacylated forms (UAGs), as listed in Table 1. These synthetic peptides all displayed the expected molecular masses, as analyzed using mass spectrometry (Table 1) and showed a single elution peak on HPLC (Fig. S4B).

### 3.2. Binding properties of fish GHSRs with the SmBiT-based tracers

To study the ligand-binding properties of these fish GHSRs, we employed the NanoBiT-based homogenous ligand-binding assay that relies on an N-terminally sLgBiT-fused receptor and a SmBiT-based tracer (Fig. 2A). Once the SmBiT-based tracer binds to the sLgBiT-fused receptor, the proximity effect would induce complementation of the ligand-attached SmBiT with the receptor-fused sLgBiT and restore NanoLuc activity. In previous studies [3,18], we prepared two SmBiT-based tracers, ghrelin-SmBiT and LEAP2-SmBiT, by chemical conjugation of a ligation version of SmBiT with a ligation version of Hs-ghrelin and Hs-LEAP2 (Fig. 2B). We first conducted saturation binding assays to test whether these fish GHSRs could bind to these tracers. When ghrelin-SmBiT or LEAP2-SmBiT were added to living HEK293T cells transiently overexpressing the sLgBiT-fused Lc-GHSR, both tracers displayed typical hyperbolic binding curves (Fig. 2C), with the calculated dissociation constants (K_d_) in the nanomolar range (Table 2), consistent with our previous data [16]. Thus, the fish GHSR from the extant coelacanth *L. chalumnae* retains high binding affinity with both tracers, although the tracers were generated based on human ghrelin and human LEAP2.

**Table 2.** Summary of the measured K_d_ values of the SmBiT-based tracers with the sLgBiT-fused GHSRs.

| Tracers | Hs-GHSR1a | Lc-GHSR1a | Dr-GHSRA | Dr-GHSRB | La-GHSR |
| --- | --- | --- | --- | --- | --- |
| Ghrelin-SmBiT | $(3.6 \pm 0.2)^a$ | $2.9 \pm 0.2$<br>$(2.7 \pm 0.2)^a$ | N.D. | N.D. | N.D. |
| LEAP2-SmBiT | $(9.5 \pm 1.3)^a$ | $4.7 \pm 0.2$<br>$(5.0 \pm 0.7)^a$ | $7.6 \pm 0.5$ | $6.1 \pm 0.2$ | $14.3 \pm 2.7$ |

<sup>a</sup> The values listed in parentheses are data from our previous study [16].
N.D., not detectable.

**Fig. 2.**
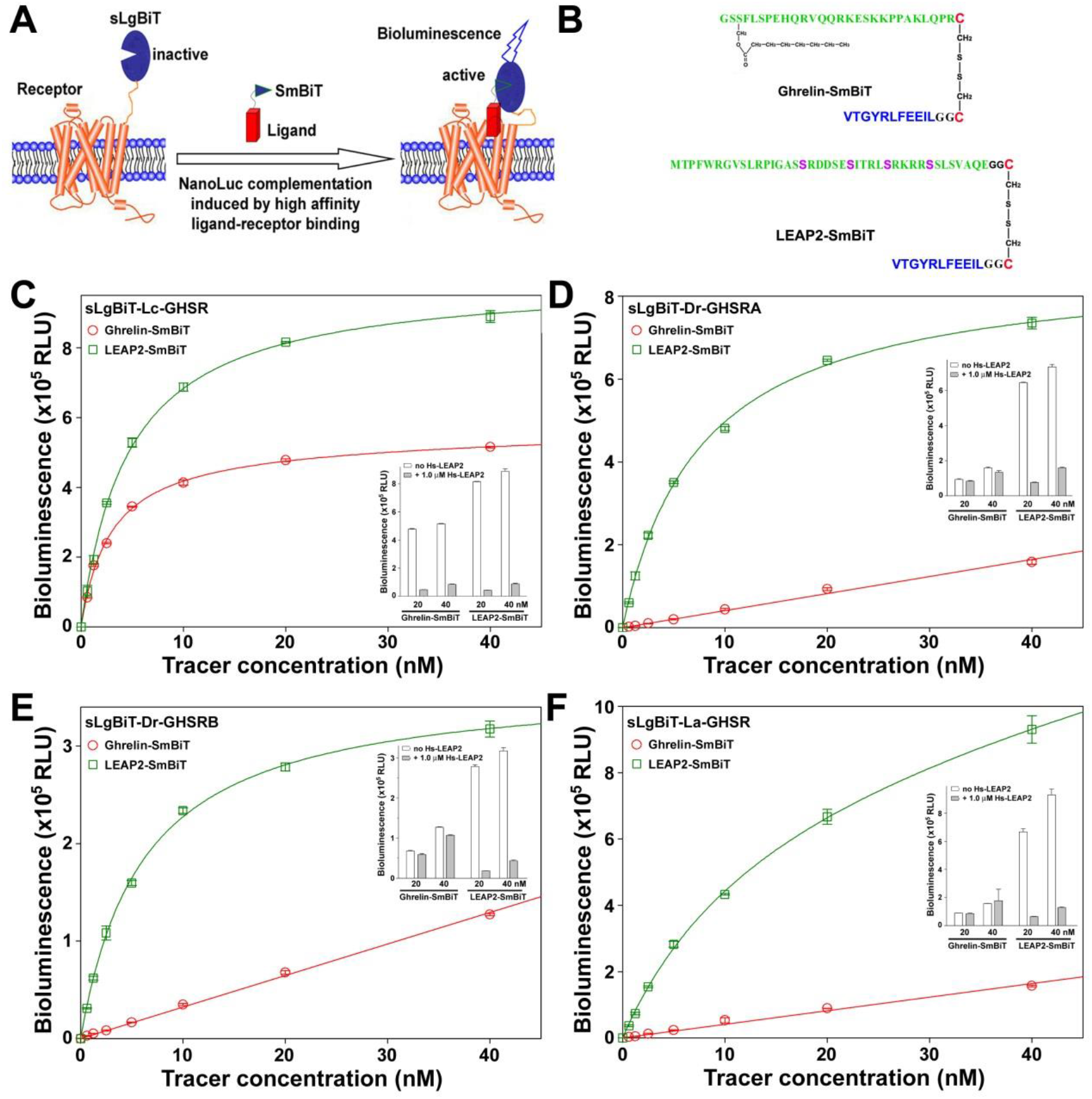
Saturation binding of the sLgBiT-fused fish GHSRs with SmBiT-based tracers. (**A**) Schematic presentation of the NanoBiT-based homogenous binding assay. (**B**) SmBiT-based tracers used in the present study. (**C-F**) Saturation binding of the sLgBiT-fused Lc-GHSR (**C**), Dr-GHSRA (**D**), Dr-GHSRB (**E**), and La-GHSR (**F**) with the SmBiT-based tracers. Inner panels, effects of Hs-LEAP2 competition on the binding of tracers with receptors.

Towards the sLgBiT-fused Dr-GHSRA and Dr-GHSRB, LEAP2-SmBiT displayed hyperbolic binding curves (Fig. 2D,E), with the calculated K_d_ values in the nanomolar range (Table 2). However, ghrelin-SmBiT displayed linear binding curves towards both receptors (Fig. 2D,E). Moreover, competition using 1.0 μM Hs-LEAP2 had only a slight effect (Fig. 2D,E, inner panels). Towards the sLgBiT-fused La-GHSR, a similar phenomenon was observed (Fig. 2F and Table 2). Thus, it seemed that the fish GHSRs from *D. rerio* and *L. crocea* retained normal binding with the LEAP2 tracer, but almost no binding with the ghrelin tracer.

### 3.3. Binding properties of fish GHSRs with unlabeled ghrelins and LEAP2s

Using LEAP2-SmBiT as a tracer, we first measured binding properties of the sLgBiT-fused GHSRs with various unlabeled ghrelins and UAGs using competition binding assays. Both Hs-GHSR and Lc-GHSR slightly preferred their cognate ghrelin, but they also efficiently bound to other ghrelins, with IC_50_ values in the nanomolar range (Fig. 3 A,B and Table 3). However, both receptors had no detectable binding with two UAGs (Table 3), suggesting that *O*-acylation is essential for ghrelin binding to Hs-GHSR and Lc-GHSR, which was consistent with previous studies [3,16].

**Table 3.** Summary of the measured IC_50_ values of LEAP2s and ghrelins binding with GHSRs and the measured EC_50_ values of ghrelins activating GHSRs.

| Peptides | IC<sub>50</sub> (nM)<sup>a</sup> or EC<sub>50</sub> (nM)<sup>b</sup> |  |  |  |  |
| --- | --- | --- | --- | --- | --- |
|  | Hs-GHSR | Lc-GHSR | Dr-GHSRA | Dr-GHSRB | La-GHSR |
| Hs-LEAP2 | 1.35 ± 0.13<br>(5.75 ± 0.56)<sup>a</sup> | 2.57 ± 0.12 | 7.59 ± 0.35 | 3.02 ± 0.14 | 3.72 ± 0.33 |
| Lc-LEAP2 | 4.79 ± 0.46 | 2.00 ± 0.09<br>(4.79 ± 0.32) | 4.37 ± 0.20 | 2.00 ± 0.09 | 2.40 ± 0.36 |
| Dr-LEAP2 | 23.4 ± 2.5 | 9.77 ± 0.65 | 5.75 ± 0.28<br>(10.7 ± 0.7) | 3.09 ± 0.14<br>(4.47 ± 0.32) | 3.80 ± 0.25 |
| La-LEAP2A | 6.17 ± 0.42 | 4.17 ± 0.19 | 2.69 ± 0.12 | 1.32 ± 0.06 | 2.14 ± 0.14<br>(15.1 ± 0.7) |
| La-LEAP2B | (>1000) | (>1000) | (>1000) | (>1000) | (>1000) |
| Hs-ghrelin | 3.63 ± 0.24<br>[2.04 ± 0.36]<sup>b</sup> | 12.3 ± 1.1<br>[4.07 ± 0.52] | (N.D.)<br>[N.D.] | (N.D.)<br>[N.D.] | (N.D.)<br>[N.D.] |
| Lc-ghrelin | 12.0 ± 0.6<br>[10.0 ± 1.5] | 6.03 ± 0.41<br>[2.57 ± 0.28] | (N.D.)<br>[N.D.] | (N.D.)<br>[N.D.] | (N.D.)<br>[N.D.] |
| Dr-ghrelin-19 | 6.92 ± 0.32<br>[4.47 ± 0.67] | 10.5 ± 0.7<br>[4.07 ± 0.52] | (>1000)<br>[N.D.] | (~1000)<br>[N.D.] | (N.D.)<br>[N.D.] |
| Dr-ghrelin-22 | 13.2 ± 0.9<br>[6.31 ± 0.94] | 11.7 ± 1.0<br>[5.50 ± 0.71] | (>1000)<br>[N.D.] | (~1000)<br>[N.D.] | (N.D.)<br>[N.D.] |
| La-ghrelin-23 | 6.61 ± 0.31<br>[5.37 ± 0.90] | 35.5 ± 1.7<br>[6.03 ± 0.41] | (N.D.)<br>[N.D.] | (~1000)<br>[N.D.] | (N.D.)<br>[N.D.] |
| Dr-UAG-22 | N.D.<br>[N.D.] | N.D.<br>[N.D.] | (N.D.)<br>[N.D.] | (N.D.)<br>[N.D.] | (N.D.)<br>[N.D.] |
| La-UAG-23 | N.D.<br>[N.D.] | N.D.<br>[N.D.] | (N.D.)<br>[N.D.] | (N.D.)<br>[N.D.] | (N.D.)<br>[N.D.] |

<sup>a</sup> The IC<sub>50</sub> values with untagged GHSRs measured via the washing-based binding assay are shown in parentheses.
<sup>b</sup> The measured EC<sub>50</sub> values for ghrelin and UAG activating GHSRs are shown in brackets.
N.D., not detectable.

**Fig. 3.**
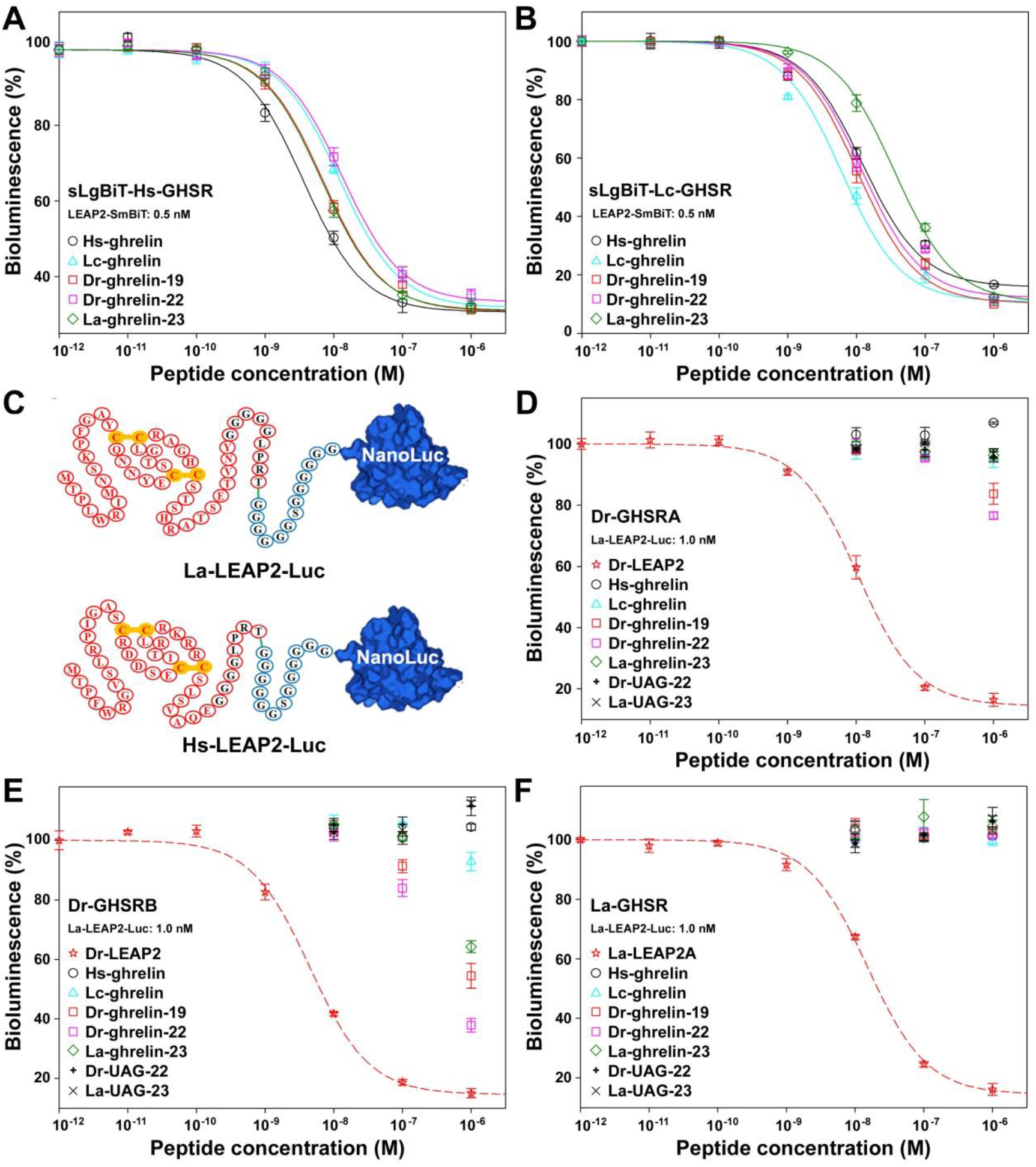
Competition binding of ghrelins and UAGs with GHSRs. (**A**,**B**) Competition binding of ghrelins with the sLgBiT-fused Hs-GHSR (**A**) and Lc-GHSR (**B**) measured via the NanoBiT-based binding assay using LEAP2-SmBiT as a tracer. (**C**) Schematic presentation of the NanoLuc-based LEAP2 tracers for the washing-based ligand-binding assays of untagged GHSRs. (**D-F**) Competition binding of ghrelins and UAGs with untagged Dr-GHSRA (**D**), Dr-GHSRB (**E**), and La-GHSR (**F**) measured via the washing-based binding assay using La-LEAP2-Luc as a tracer.

Unfortunately, the sLgBiT-fused Dr-GHSRA, Dr-GHSRB, and La-GHSR showed no detectable binding with all ghrelins and UAGs in the NanoBiT-based competition binding assays. To confirm this result, we conducted washing-based binding assays on untagged receptors using the newly developed NanoLuc-based LEAP2 tracers that were generated using sortase A-catalyzed ligation of Hs-LEAP2-srt and La-LEAP2-srt with 6×Gly-NanoLuc (Fig. 3C). In the washing-based competition binding assays, Dr-GHSRA, Dr-GHSRB, and La-GHSR all bound to their cognate LEAP2 with a typical sigmoidal curve (Fig. 4E−F) and the calculated IC_50_ values were comparable to those measured using the NanoBiT-based binding assays (Table 3). However, the three untagged GHSRs showed almost no binding to all ghrelins and UAGs (Fig. 4E−F and Table 3), consistent with the fact that they showed no binding with the ghrelin-SmBiT tracer in the NanoBiT-based saturation binding assays (Table 2).

**Fig. 4.**
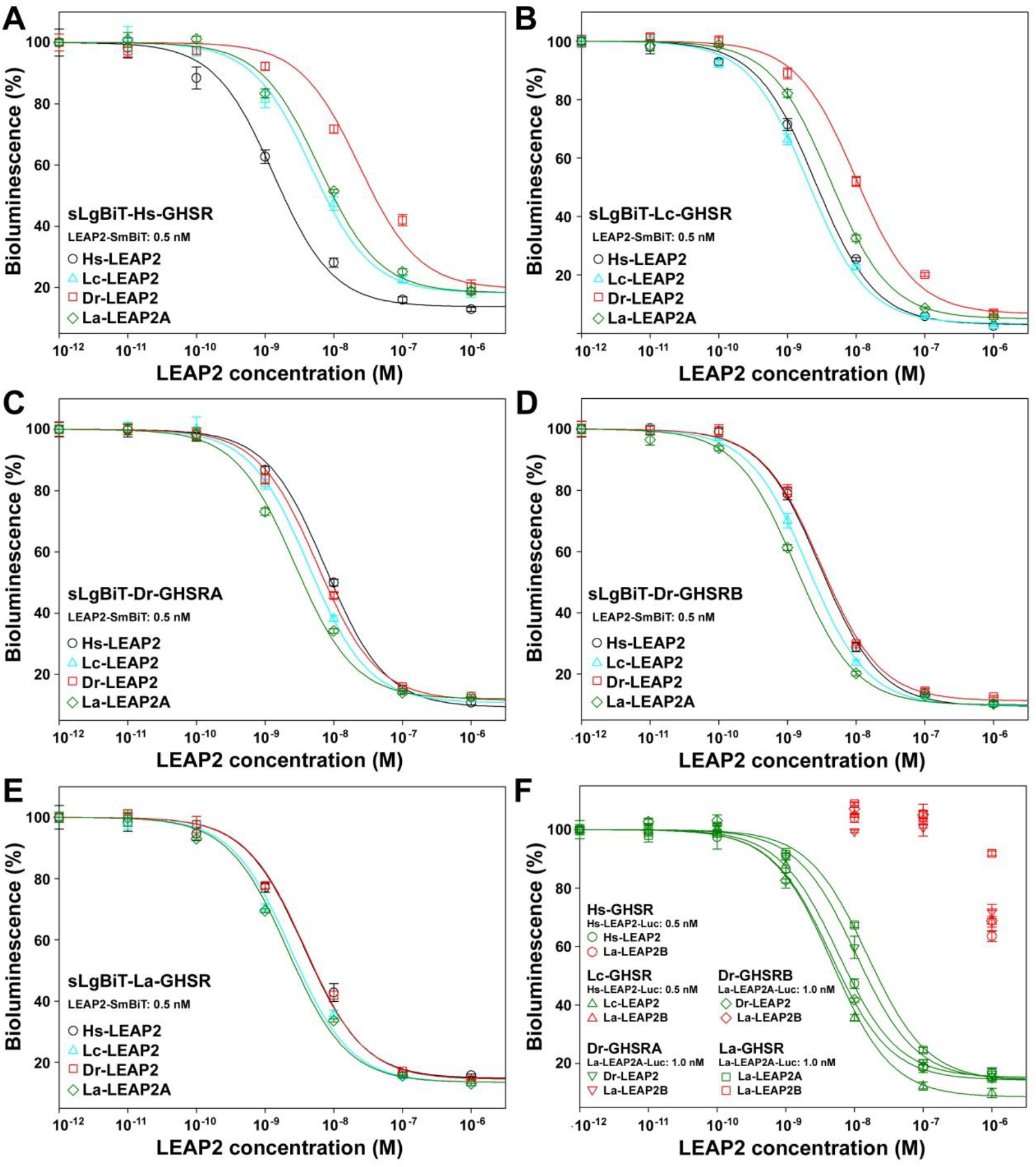
Competition binding of LEAP2s with GHSRs. (**A-E**) Competition binding of active LEAP2s with the sLgBiT-fused Hs-GHSR (**A**), Lc-GHSR (**B**), Dr-GHSRA (**C**), Dr-GHSRB (**D**), and La-GHSR (**E**) measured via the NanoBiT-based binding assay using LEAP2-SmBiT as a tracer. (**F**) Competition binding of La-LEAP2B with untagged GHSRs measured via the washing-based binding assay.

Subsequently, we measured the binding properties of the sLgBiT-fused GHSRs with various unlabeled LEAP2s using competition binding assays (Fig. 4A-E). All GHSRs displayed almost no detectable binding with La-LEAP2B; however, they bound to other LEAP2s tightly, with the calculated IC_50_ values in the nanomolar range (Table 3). Hs-GHSR markedly preferred its cognate ligand Hs-LEAP2 (Fig. 4A and Table 3), whereas the fish GHSRs displayed similar binding potencies towards all active LEAP2s (Fig. 4B-E and Table 3). Thus, it seemed that these GHSRs all have a functional LEAP2 as an endogenous ligand, and displayed high cross activity with LEAP2s from other species.

To confirm the low activity of La-LEAP2B, we conducted the washing-based ligand-binding assays on untagged GHSRs using the NanoLuc-based LEAP2 tracers as shown in Fig. 3C. In the washing-based binding assays, all active LEAP2s displayed typical sigmoidal competition curves towards their cognate receptors, with calculated IC_50_ values comparable to those measured using the NanoBiT-based binding assays (Table 3). In contrast, the binding activity of La-LEAP2B towards all GHSRs was too low to be accurately quantified (Fig. 4F and Table 3). Thus, it seemed that La-LEAP2B has lost almost all its binding activity not only to its cognate receptor, but also to GHSRs from other species. Fortunately, La-LEAP2A is highly active, thus *L. crocea* has a functional LEAP2 as an endogenous ligand of its GHSR.

### 3.3. Activation of GHSRs by ghrelins

We measured the activation potencies of ghrelins towards GHSRs via a gene receptor assay using a CRE-controlled NanoLuc reporter. Although GHSRs signal through the Gq pathway, they also activate the transcription factor CREB through kinases such as Ca^2+^/calmodulin kinase IV and protein kinase C [22]. Thus, the CRE-controlled luciferase reporter assay can be used to monitor GHSR activation, as demonstrated in previous studies [3,16−18,23]. Hs-GHSR and Lc-GHSR were efficiently activated by all synthetic octanoylated ghrelins (Fig. 5A), with calculated EC_50_ values in the nanomolar range (Table 3), consistent with the fact that they could bind tightly to all the tested ghrelins. In contrast, we detected no activation of Hs-GHSR and Lc-GHSR by unacylated UAGs (Fig. 5A), consistent with the fact that they had no detectable binding with both receptor (Table 3). For the GHSRs from *D. rerio* and *L. crocea*, all synthetic ghrelins and UAGs had no detectable activation effect (Table 3), consistent with the fact that these receptors showed almost no binding to any of the tested ghrelins.

**Fig. 5.**
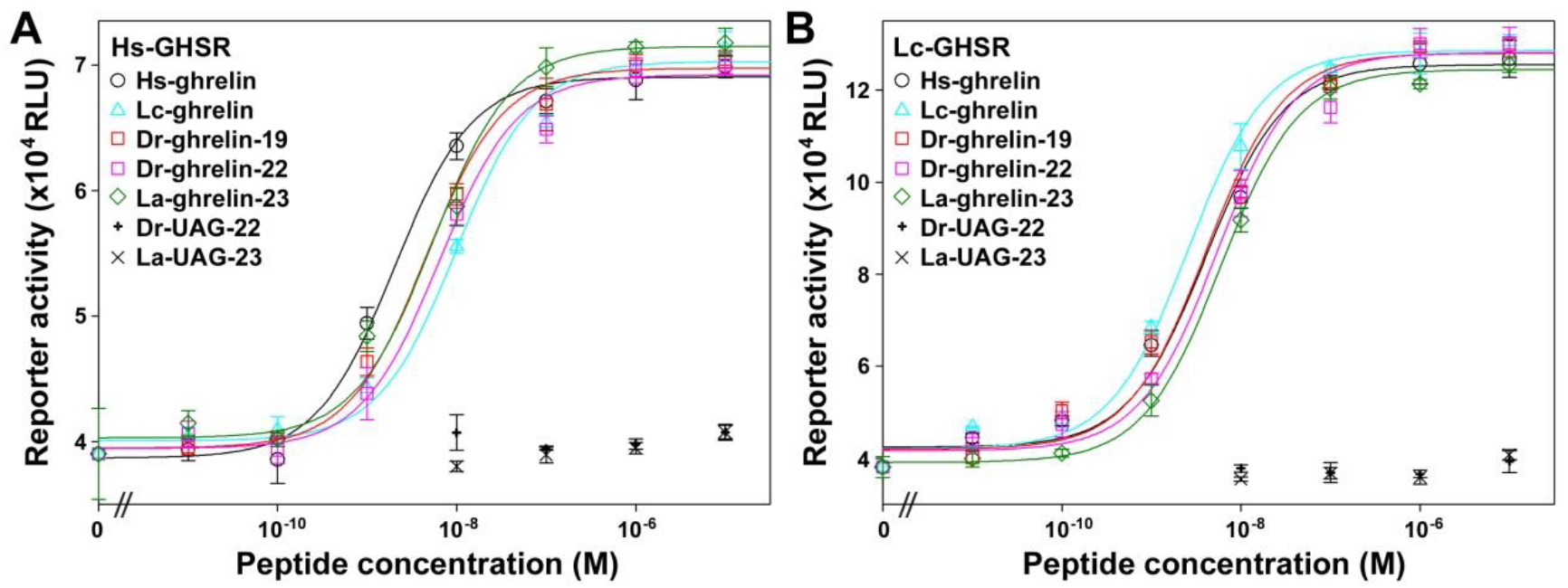
Activation of untagged Hs-GHSR (**A**) and Lc-GHSR (**B**) by synthetic ghrelins and UAGs monitored using a CRE-controlled NanoLuc reporter.

## 4. Discussion

The present study demonstrated that both *D. rerio* and *L. crocea* have a functional LEAP2 that not only binds to their cognate GHSRs with nanomolar range affinity, but also displays high cross reactivity with human and coelacanth GHSRs. Thus, it seems that LEAP2 functions as a ligand of GHSR in all fish and this function is highly conserved in evolution from fish to mammals. Surprisingly, all synthetic n-octanoylated ghrelins showed almost no binding and activation effects on GHSRs from *D. rerio* and *L. crocea*; however, these peptides displayed normal functions towards human and coelacanth GHSRs. Thus, it seemed that GHSRs from *D. rerio* and *L. crocea* have lost their response to ghrelin. Although GHSRs have been identified from many fish species, only a few of them have been functionally characterized [24−30]. As summarized in Table 4, those fish GHSRs carrying an aromatic Phe residue at their 6.58 position are generally active towards ghrelins; however, other fish GHSRs generally display low activity or even no activity towards ghrelins. It is somewhat confusing that zGHS-R1a (corresponding to Dr-GHSRA) was reported as active towards goldfish ghrelin, but inactive towards rat ghrelin [24], considering the high sequence similarity of the two orthologs [31]. This discrepancy needs to be clarified in future studies. Without ghrelin as an agonist, LEAP2 might function as an inverse agonist of GHSR in these fish; however, more studies need to be carried out in the future. These functional assays were conducted in transfected mammalian cells; therefore, we cannot exclude the possibility that the overexpressed fish receptors had some defects and thus lost their response to ghrelins. However, this possibility was not high considering that those fish GHSRs carrying a Phe residue at their 6.58 position are generally active towards ghrelins, and all hitherto tested fish GHSRs display normal binding with LEAP2s.

**Table 4.** Summary of the functional characterization of fish GHSRs in the present and previous studies.

| Fish species | GHSRs | Residue<br>at 6.58 | Assays and results | References |
| --- | --- | --- | --- | --- |
| *Danio rerio* | Dr-GHSRA | Ile | Binding assay: almost lost binding to ghrelins, but retained normal binding to LEAP2s. | present study |
|  | or |  | CRE-reporter: no response to all tested ghrelins. |  |
|  | zGHS-R1a |  | Calcium mobilization: no response to rat ghrelin, but active to goldfish ghrelin. | 24 |
|  | Dr-GHSRB | Val | Binding assay: lost most binding to ghrelins, but retained normal binding to LEAP2s. | present study |
|  | or |  | CRE-reporter: no response to all tested ghrelins. |  |
|  | zGHS-R2a |  | Calcium mobilization: no response to rat ghrelin and goldfish ghrelin. | 24 |
| *Oncorhynchus mykiss* | rtGHSR1a-LR | Met | Calcium mobilization: no response to rat ghrelin and des-VRQ trout ghrelin. | 25 |
| *Carassius auratus* | GHS-R1a-1 | Ile | Calcium mobilization: low activity to goldfish ghrelin and rat ghrelin. | 26 |
|  | GHS-R1a-2 |  | Calcium mobilization: low activity to goldfish ghrelin, but no activity to rat ghrelin. |  |
|  | GHS-R2a-1 | Phe | Calcium mobilization: active to goldfish ghrelin and rat ghrelin. |  |
|  | GHS-R2a-2 |  | Calcium mobilization: no activity. |  |
| *Larimichthys crocea* | La-GHSR | Gln | Binding assay: lost binding to ghrelins, but retained normal binding to LEAP2s. | present study |
|  |  |  | CRE-reporter: no response to all tested ghrelins. |  |
| *Acanthopagrus schlegeli* | sbGHSR-1a | Gln | Calcium mobilization: low activity stimulated by 10 μM human ghrelin. | 27 |
| *Oreochromis mossambicus* | GHS-R1a | Gln | Calcium mobilization: no response to tilapia or rat ghrelins. | 28 |
| *Latimeria chalumnae* | Lc-GHSR | Phe | Binding assay: high affinity binding to ghrelins and LEAP2s. | Present study, 16 |
|  |  |  | CRE-reporter: efficient activation by ghrelins. |  |
| *Ictalurus punctatus* | GHS-R1a | Phe | Only GHS-R3a was functionally characterized by calcium mobilization: active to catfish ghrelin. | 29 |
|  | GHS-R2a |  |  |  |
|  | GHS-R3a |  |  |  |
| *Protopterus annectens* | GHSR1a | Phe | Calcium mobilization: active to rat ghrelin and bullfrog ghrelin. | 30 |

Recently, high resolution structures of human GHSR complexed with ghrelin and G-proteins have been solved by two laboratories using cryo-electron microscopy [32,33]. These structures suggested that the conserved Phe286 (Phe6.58) of human GHSR interacts with the hydrophobic Leu5 of ghrelin. This could explain why those fish GHSRs without a Phe residue at their 6.58 position have generally lost their binding to ghrelins. However, the importance of Leu5 for ghrelin function was underestimated for a long time, because an early study mistakenly reported that Ala replacement of Leu5 had no detrimental effect on ghrelin function [34]. Recently, we demonstrated that Leu5 is actually important for ghrelin to bind to and activate GHSR [17]. Leu5 is highly conserved in ghrelins from fish to mammals, except for two orthologs from *Cavia porcellus* (guinea pig) and *Phyllostomus discolor* (pale spear-nosed bat), which bear concurrent genetic variations, Ser2 to Ala and Leu5 to Arg, compared with other ghrelins. However, the unusual ghrelin ortholog from *C. porcellus* has been demonstrated to be fully active [35]. Our recent study suggested that the positively charged Arg5 in the unusual ghrelin orthologs likely forms cation−π and π−π interactions with Phe6.58 of GHSR, and thus compensates for the loss of the hydrophobic Leu5 residue [17].

Quick progress in DNA sequencing technologies has led to more and more protein orthologs from various species becoming available, thus “how to use these sequences” has become a question. All known GHSRs from fish to mammals share high overall sequence similarity, implying that they have conserved functions. However, we noticed that most fish GHSRs lack an aromatic Phe residue at their 6.58 position. This residue is highly conserved from amphibians to mammals, and has been demonstrated to be important for human GHSR binding to ghrelin, although it is irrelevant for the receptor binding to LEAP2 [18,19]. We originally speculated that these unusual fish GHSRs might have evolved an alternative mechanism for efficient binding to ghrelin, similar to the unusual ghrelin orthologs from *C. porcellus* and *P. discolor* [17]. However, this speculation was not supported by our present results and some previous studies. It seemed that these fish GHSRs really have lost their response to ghrelins, although they retain normal binding to LEAP2s. In future studies, the functions of the ghrelin-LEAP2-GHSR system in different fish species should be further investigated.

## Acknowledgments

This work was supported by grant from the National Natural Science Foundation of China (31971193).

