## Supplementary Fig. S1-S5 for "LEAP2 is a more conserved ligand than ghrelin for fish GHSRs"

**Contents:**

**Fig. S1.** Amino acid sequence alignment of fish GHSRs using human GHSR as a control.

**Fig. S2.** The nucleotide and amino acid sequences of LEAP2 precursors overexpressed in *E. coli*.

**Fig. S3.** The nucleotide and amino acid sequence of 6×His-6×Gly-NanoLuc overexpressed in *E. coli*.

**Fig. S4.** HPLC analyses of the synthetic ghrelins (A) and recombinant LEAP2s (B).

**Fig. S5.** Amino acid sequence alignment of MBOAT4 (GOAT) from *Homo sapiens*, *Latimeria chalumnae*, *Danio rerio*, and *Larimichthys crocea*.

### Alignment of Fish GHSRs using human GHSR as a control

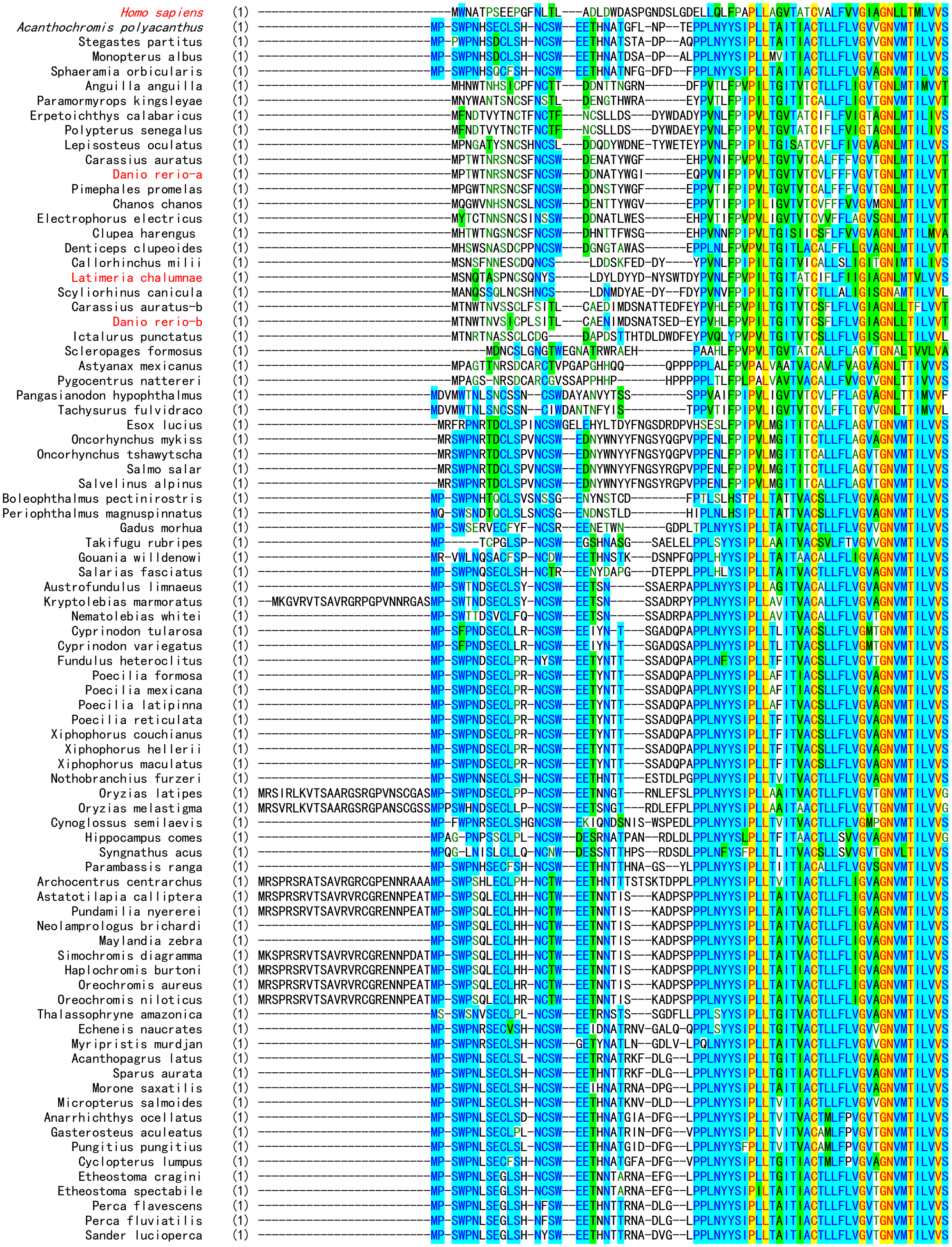

```
Cottoperca gobio                    (1)  -----------------------------MMGDRSVCLSH-NCSL--EETHNATRNA-DLG--KPPLYYYTIPLLTVITIACTLLFLVGVTGNVMTILVVS
Gymnodraco acuticeps                (1)  --------------------------MP-SWPNLSEFLSH-NCCW--EETINATRTA-EL----PPLNYYSIPLLTGITIACTLLFLVGVTGNVMTMLVVS
Pseudochaenichthys georgianus       (1)  --------------------------MP-SWPNLSEFLSH-NCCW--EETINATRTA-EL----PPLNYYSIPLLTGITIACTLLFLVGVTGNVMTMLVVS
Notothenia coriiceps                (1)  --------------------------MP-SWPNLSEFLSH-NCCW--EETINATGTA-EQ----PPLNYYSIPLLTGITIACTLLFLVGVTGNVMTMLVVS
Trematomus bernacchii               (1)  --------------------------MP-SWPNLSEFLSR-NCCW--EEIINATSTA-EL----PPLNYYSIPLLTGITIACTLLFLVGVTGNVMTMLVVS
Epinephelus lanceolatus             (1)  --------------------------MP-SWPNLSLCLSH-NCSW--EETHNATRNA-DLG--LPPLNYYSIPLLTVITVACTLLFLIGVTGNVMTILVVS
Larimichthys crocea                 (1)  --------------------------MP-SWPNLSECLSL-NCSW--EETDNVTRNA-DLG--RPPTNYFSIPLLTATTIACTLLFLVGVVGNVMTILVVS
Sebastes umbrosus                   (1)  --------------------------MP-SCPNLSECLSHDNYSW--EETHNNATRT-SD-LGLPPLNYYSIPLLTAITTACTLLFLVGVTGNVMTILVVG
Anabas testudineus                  (1)  --------------------------MP-SWPNHSECLSH-NCSG--EENHNAT------DHVLPPLNYYSIPLLTAITIACTLLFLVGVAGNVMTILVVS
Betta splendens                     (1)  --------------------------MP-SLPNRSECVSH-SC-W--EEKHNATRD---ADPVVPPLNYYSIPLLTGITIACTLLFLVGVAGNVMTILVVS
Hippoglossus hippoglossus           (1)  --------------------------MP-TWPNRSECVSH-NCSW--EKIPNATWK---DDYVLPPLNYYSIPLLTAITTACTLLFLVGMAGNVMTILVVS
Hippoglossus stenolepis             (1)  -MSPPPGVPSTLRARRGPVNNRGDSMP-TWPNRSECVSH-NCSW--EKIPNATWK---DDYVLPPLNYYSIPLLTAITTACTLLFLVGMAGNVMTILVVS
Paralichthys olivaceus              (1)  --------------------------MP-TWPNHSECFSH-NCSW--EKIHNATWN---DDYVLPPLNYYSIPLLTAITVACTLLFLVGMAGNVMTILVVS
Scophthalmus maximus                (1)  --------------------------MP-TWPNHSECVSH-NCSG--EEIHNSTWD---DDHVLPPLNYYSIPLLVVITVACTLLFLVGMAGNVMTILVVS
Labrus bergylta                     (1)  --------------------------MP-PWQNQSQCISQ-NCSW--ETLNSTGDTD-LG---WPPLNYYSIPLLTAITIACTLLFLVGVAGNVMTILVVS
Notolabrus celidotus                (1)  --------------------------MP-SLPNQSQCLSR-NCSW--EEIHNN-TAD-LG---LPPLNYYSIPLLTAITIACTLLFLVGVVGNVMTILVVS
Seriola dumerili                    (1)  --------------------------MP-SWLNHSECISH-NCSW--EEINDTTRN---DDPVLPPLNYYSIPLLTGITIACTLLFLVGMTGNVMTILVVS
Seriola lalandi dorsalis            (1)  --------------------------MP-SWLNHSECISH-NCSW--EEINDTTRN---DDPVLPPLNYYSIPLLTGITIACTLLFLVGMTGNVMTILVVS
Toxotes jaculatrix                  (1)  --------------------------MP-SWPNHSECLSH-NCSW--EEIHNTTRN---DDPVLPPLNYYSIPLLTAITIACTVLFLVGVVGNVMTILVVS
Lates calcarifer                    (1)  --------------------------MP-SWPNHSECLSH-NCSW--EEIHNHTRN---ADPVPPPLNYYSIPLLTAITIACTLLFLVGVAGNVMTILVVS
Xiphias gladius                     (1)  --------------------------MP-SWPNHSECLSH-NCSW--GEIHNATRN---ADPILPPLNYYSIPLLTAITIACTLLFLVGVAGNVMTILVVS
Mastacembelus armatus               (1)  --------------------------MP-SWPNHSECLSH-NCSW--DETNNATRN---DDPVLPPLNYYSIPLLTAITIACTLLFLVGVAGNVMTILVVS

Homo sapiens                        (70) RFRELRTTTNLYLSSMAFSDLLIFLCMPLDLVRLWQYRPWNFGDLLCKLFQFVSESCTYATVLTITALSVERYFAICFPLRAKVVVTKGRVKLVIFVIWA
Acanthochromis polyacanthus         (69) KYRDMRTTTNLYLCSMAVSDLLIFLCMPLDLYRMWRYRPWRFGDALCKLFQFVSESCTYSTILSITALSVERYLAICFPLRAKALVTKRRVRALILLLWT
Stegastes partitus                  (69) KYRDMRTTTNLYLCSMAVSDLLIFLCMPLDLYRMWRYRPWRFGDALCKLFQFVSESCTYSTILSITALSVERYLAICFPLRAKALVTKRRVRALILLLWT
Monopterus albus                    (69) KYRDMRTTTNLYLCSMAVSDLLIFLCMPLDLYRMWRYRPWRFGDALCKLFQFVSESCTYSTILSITALSVERYLAICFPLRAKALVTKRRVRALILLLWT
Sphaeramia orbicularis              (69) KYRDMRTTTNLYLCSMAVSDLLIFVCMPLDLYRMWRYRPWRFGDALCKLFQFVSESCTYSTILSITALSVERYLAICFPLRAKALVTKRRVRALILLLWT
Anguilla anguilla                   (64) KYKDMRTTTNLYLSSMALSDLLIFLCMPLDLYRVWRYRPWNFGDLLCKLFQFVSESCTYSTILNITALSVERYFAICFPLRAKVVVTKGRVRGVILVLWM
Paramormyrops kingsleyae            (64) MYKDMRTTTNLYLSSMALSDLLIFICMPLDLYRIWRYRPWNFGDLLCKLFQFISESCTYSTILSITALSVERYFAICFPLRAKVLVTKGRVRGIILLLWI
Erpetoichthys calabaricus           (67) KYKDMRTTTNLYLSSMAFSDLFIFLCLPLDLYRIWRYRPWNFGDLLCKLFQFVSESCTYSTILNITALSIERYFAICFPLRAKVVVTKGRVKGIILLLWA
Polypterus senegalus                (67) KYKDMRTTTNLYLSSMAFSDLFIFLCLPLDLYRIWRYRPWNFGDLLCKLFQFVSESCTYSTILNITALSIERYFAICFPLRAKVVVTKGRVKGIILLLWA
Lepisosteus oculatus                (69) KFKDMRTTTNLYLSSMAFSDLLIFLCMPLDLYRIWRYRPWNFGDLLCKLFQFISESCTYSTILNITALSVERYFAICFPLRAKVVVTKSRVKGIILVLWL
Carassius auratus                   (64) KYKDMRTTTNLYLSSMAFSDLLIFLCMPLDLYRIWRYRPWNFGNVLCKLFQFVSECCTYSTILNITALSVERYFAICFPLRAKVVVTKGRVRGVILVLWI
Danio rerio-a                       (64) KYKDMRTTTNLYLSSMAFSDLLIFLCMPLDLYRIWRYRPWNFGNILCKLFQFVSECCTYSTILNITALSVERYFAICFPLRAKVVVTKGRVRGVILVLWI
Pimephales promelas                 (64) KYKDMRTTTNLYLSSMAFSDLLIFLCMPLDLYRLWKYRPWNFGNILCKLFQFVSECCTYATILNITALSVERYFAICFPLRAKIVVTKGRVRGVILVLWI
Chanos chanos                       (64) RYKDMRTTTNLYLSSMAFSDLLIFLCMPLDLYRIWRYRPWNLGDILCKLFQFVSECCTYSTILNITALSVERYFAICFPLRAKVVVTKGRVRGVILVLWI
Electrophorus electricus            (64) KYKDMRTTTNLYLSSMAFSDLLIFLCMPLDLYRIWRYRPWNFGNALCKLFQFISECCTYSTILNITALSVERYFAICFPLRAKVVVTKGRVRGVILFLWT
Clupea harengus                     (64) KYRDMRTTTNLYLSSMALSDLLIFLCMPLDLYRVWRYRPWYFGDLLCKLFQFVSEGCTYSTILNITALSVERYFAICFPFRAKVVVTKGRVRGIILILWV
Denticeps clupeoides                (64) RYKDMRTTTNLYLSSMALSDLLIFLCMPLDLYRIWRYRPWNFGDALCKLFQFISECCTYSTILNITALSVERYFAICFPLRAKVVVTKGRVRGIILLLWA
Callorhinchus milii                 (63) KFRDMRTTINFYLSSMALSDLMIFMCMPFDLYRIWQYRPWNFGVMLCKMFQFISESCTYSTILNITALSVERYLAICFPLKAKVVITKGRVKGIILFLWT
Latimeria chalumnae                 (67) KFKEMRTTTNLYLSSMAFSDLLIFLCMPLDLFRIWQYRPWNFGDLLCKLFQFVSESCTYSTILSITALSVERYFAICFPLKAKVIITKGRVKLVILLLWV
Scyliorhinus canicula               (65) KFKGMRTTTNFYLSSMAFSDLFIFVCMPLDLYRIWQYRPWNFGSLLCKLFQFVSESCTYSTILNITALSVERYFAICFPLKAKVVITKGRVKGLIFFLWS
Carassius auratus-b                 (70) KYKDMRTTTNLYLCSMALSDLLIFLCMPLDLYRVWRYRPWNFGDELCKLFQFVSESCTYSTILNITALSVERYFAICFPFRAKVIVTRGRVKGVILLLWT
Danio rerio-b                       (69) KYKDMRTTTNLYLCSMALSDLLIFLCMPLDLYRVWRYRPWNFGDELCKLFQFVSESCTYSTILNITALSVERYFAICFPLRAKVIVTRGRVKGVILLLWT
Ictalurus punctatus                 (68) KYKDMRTTTNLYLCSMALSDLLIFLCMPLDLYRMWRYRPWNLGDHLCKLFQFISESCTYSAVLSITALSVERYVAICFPLRAKVLVTRGRVKALILLLWL
Scleropages formosus                (60) KFKDMRTTTNLYLCSMAVSDLLVLLWLPLDLYRVWRYRPWGFGQLLCKLFQFVSESCTYSSILSITALSVERYLAICFPLRAKVLVTRARVRGLLAALWL
Astyanax mexicanus                  (67) RYRDMRTTTNLYLRSMALSDLLVFACMPLDLQRVWRYRPWPLGGAVCKLFHFVSESCTHCTALSIAALSVHRYLAVRFPLHAKVAVTRRRVRAVILALWA
Pygocentrus nattereri               (65) RYRDMRTTTNLYLRSMALSDLLVFACMPLDLQRVWRYRPWPLGGAVCKLFHFVSECCTHCTALSIAALSVHRYLAVRFPLRAKVSVTRRRVRAVILALWA
Pangasianodon hypophthalmus         (66) KYKEMRTTTNLYLSSMAFSDLLIFLCLPLDLYRMWRYRPWTLGNALCKLFQFVSECCTYSTILNMTALSAERYFAICFPLRAKVAVTKRRVRGLILALWT
Tachysurus fulvidraco               (65) KYKEMRTTTNLYLSSMAFSDLLIFLCLPLDLYRMWRYRPWMLGNVLCKLFQFVSECCTYSTILNMTALSAERYFAICFPLRAKVTVTKRRVRGLILSLWT
Esox lucius                         (75) KYRDMRTTTNLYLCSMAVSDLLIFLCMPPDLYRLWKYRPWRFGDTFCKLFQFVSECCTYSTILNITALSVERYLAICFPLRAKRLVTKRRVRALILFLWL
Oncorhynchus mykiss                 (73) KYRDMRTTTNLYLCSMAVSDLLIFLCMPPDVYRLWKYRPWIFGDTFCKLFQFVSECCTYSTILNITALSVERYLAICFPLRAKRLVTKRRVRALILFLWL
Oncorhynchus tshawytscha            (73) KYRDMRTTTNLYLCSMAVSDLLIFLCMPPDVYRLWKYRPWIFGDTFCKLFQFVSECCTYSTILNITALSVERYLAICFPLRAKRLVTKRRVRALIIFLWL
Salmo salar                         (73) KYRDMRTTTNLYLCSMAVSDLLIFLCMPPDVYRLWKYRPWIFGDTFCKLFQFVSECCTYSTILNITALSVERYLAICFPLRAKRLVTKRRVRALILFLWL
Salvelinus alpinus                  (73) KYRDMRTTTNLYLCSMAVSDLLIFLCMPPDVYRLWKYRPWIFGDTFCKLFQFVSECCTYSTILNITALSVERYLAICFPLRAKRLVTKRRVRALILFLWL
Boleophthalmus pectinirostris       (65) KYRDMRTTTNLYLCSMAVSDLVILLCMPLDLYRMWRYRPWRFGDALCKLFNFVSELCTYSTILSITALSVERYLAICFPLRAKALVTKKRVRALILVLWT
Periophthalmus magnuspinnatus       (65) KFRDMRTTTNLYLCSMAVSDLVILLCMPLDLYRMWRYRPWRFGDALCKLFNFVSELCTYSTILSITALSVERYLAICFPLRAKALVTKKRVRALILALWT
Gadus morhua                        (66) KYRDMHTTTNLYLCSMAVSDLLIFLCMPPDLYRMWRYRPWRFGDALCKLFQFVSESCTYSTILSITALSVERYLAICFPLRAKALVTKRRVRALILLLWI
Takifugu rubripes                   (65) RYRDMRTTTNLYLCSMAVSDLFIFVCMPLDLYRMWRYRPWRFGNALCKLFQFVSESCTYSTILCITALSVERYLAICFPLRAKALVTKRRVRALILLLWT
Gouania willdenowi                  (69) KYRDMRTTTNLYLCSMAVSDLLIFLCMPLDLYRMWRYRPWRFGVALCKLFQFVSESCTYSTILSITALSVERYLAICFPLRVKALVTKRRVRALILFLWT
Salarias fasciatus                  (69) KYRDMRTTTNLYLCSMAVSDLLIFFCMPLDLYRMWNYRPWRFGDALCKLFQFVSESCTYSTILSITALSVERYLAICFPLRAKALVTKKRVRALILFLWT
Austrofundulus limnaeus             (67) KYRDMRTTTNLYLCSMAVSDLLIFFCMPLDLYRMWRYRPWRFGDALCKLFQFVSESCTYSTILNITALSVERYLAICYPLRAKALVTKRRVRALIFILWT
Kryptolebias marmoratus             (90) KYRDMRTTTNLYLCSMAVSDLLIFLCMPLDLYRMWRYRPWRFGDALCKLFQFVSESCTYSTILNITALSVERYLAICFPLRAKALVTKRRVRALIFIIWT
Nematolebias whitei                 (67) KYRDMRTTTNLYLCSMAVSDLLIFLCMPLDLYRMWRYRPWRFGDALCKLFQFVSESCTYSTILNITALSVERYLAICFPLRAKALVTKRRVRALIFILWT
Cyprinodon tularosa                 (68) KYRDMRTTTNLYLCSMAMSDLLIFLCMPLDLYRMWRYRPWNLGDALCKLFQFVSESCTYSTILNITALSVERYLAICFPLRAKALVTKRRVRALICILWT
Cyprinodon variegatus               (68) KYRDMRTTTNLYLCSMAVSDLLIFLCMPLDLYRMWRYRPWNLGDALCKLFQFVSESCTYSTILNITALSVERYLAICFPLRAKALVTKRRVRALICILWT
Fundulus heteroclitus               (69) KYRDMRTTTNLYLCSMAVSDLLIFFCMPLDLYRMWRYRPWRLGDALCKLFQFVSESCTYSTILNITALSVERYLAICFPLRAKALVTKRRVRALICILWT
Poecilia formosa                    (69) KYRDMRTTTNLYLCSMAVSDLLIFLCMPLDLYRMWRYRPWRLGDALCKLFQFVSESCTYSTILNITALSVERYLAICFPLRAKALVTKRRVRALICVLWT
Poecilia mexicana                   (69) KYRDMRTTTNLYLCSMAVSDLLIFLCMPLDLYRMWRYRPWRLGDALCKLFQFVSESCTYSTILNITALSVERYLAICFPLRAKALVTKRRVRALICVLWT
Poecilia latipinna                  (69) KYRDMRTTTNLYLCSMAVSDLLIFLCMPLDLYRMWRYRPWRLGDALCKLFQFVSESCTYSTILNITALSVERYLAICFPLRAKALVTKRRVRALICVLWT
Poecilia reticulata                 (69) KYRDMRTTTNLYLCSMAVSDLLIFLCMPLDLYRMWRYRPWRLGAALCKLFQFVSESCTYSTILNITALSVERYLAICFPLRAKALVTKRRVRALICVLWT
Xiphophorus couchianus              (69) KYRDMRTTTNLYLCSMAVSDLLIFLCMPLDLYRMWRYRPWRLGDALCKLFQFVSESCTYSTILNITALSVERYLAICFPLRAKALVTKRRVRALICVLWT
Xiphophorus hellerii                (69) KYRDMRTTTNLYLCSMAVSDLLIFLCMPLDLYRMWRYRPWRLGDALCKLFQFVSESCTYSTILNITALSVERYLAICFPLRAKALVTKGRVRALICILWT
Xiphophorus maculatus               (69) KYRDMRTTTNLYLCSMAVSDLLIFLCMPLDLYRMWRYRPWRLGDALCKLFQFVSESCTYSTILNITALSVERYLAICFPLRAKALVTKRRVRALICVLWT
Nothobranchius furzeri              (69) RFRDMRTTTNLYLCSMAVSDLLIFLCMPLDLYRMWRYRPWRFGDALCKLFQFVSESCTYSTILNITALSVERYLAVCFPLRAKALVTKKRVRALIFVLWT
Oryzias latipes                     (94) RYRDMRTTTNLYLCSMAVSDLLIFLCMPLDLYRMWRYRPWRLGDALCKLFQFVSESCTYSTILSITALSVERYLAICFPLRAKALVTKRRVRALICFLWT
Oryzias melastigma                  (95) RYRDMRTTTNLYLCSMAVSDLLIFLCMPLDLYRMWRYRPWRLGDALCKLFQFVSESCTYSTILSITALSVERYLAICFPLRAKALVTKRRVRALIFFLWT
Cynoglossus semilaevis              (72) KYRDMRTTTNLYLCSMAVSDLLIFLCMPLDLYRMWRYRPWLLGDALCKLFQFVSESCTYSTILSITALSVERYLAICFPLRAKALVTKRRVRALILLLWT
Hippocampus comes                   (70) KYRDMRTTTNLYLCSMAVSDLLIFLCMPLDLYRMWRYRPWRLGDALCKLFQFVSESSTYSTILSITALSVERYVAICFPLRAKALVTKRRVRALILLLWC
Syngnathus acus                     (70) KYRDMRTTTNLYLCSMAVSDLLIFLCMPLDLYRMWRYRPWRLGDALCKLFQFVSESSTYSTILSITALSVERYVAICIPLRAKALMTKRRVRSLILLLWC
Parambassis ranga                   (69) KYRDMRTTTNLYLCSMAVSDLLIFLCMPLDLYRMWRYRPWRLGDVLCKLFQFVSESCTYSTILSITALSVERYMAICFPLRAKALVTKRRIRALILFLWT
Archocentrus centrarchus            (97) KYRDMRTTTNLYLCSMAVSDLLIFLCMPLDLYRMWRYRPWRFGDALCKLFQFVSESSTYSTILSITALSVERYLAICFPLRAKALVTKRRVRALICLLWT
Astatotilapia calliptera            (94) KYRDMRTTTNLYLCSMAVSDLLIFLCMPLDLYRMWRYRPWRFGDALCKLFQFVSESSTYSTILSITALSVERYLAICFPLRAKALVTKRRVRALICLLWT
Pundamilia nyererei                 (94) KYRDMRTTTNLYLCSMAVSDLLIFLCMPLDLYRMWRYRPWRFGDALCKLFQFVSESSTYSTILSITALSVERYLAICFPLRAKALVTKRRVRALICLLWT
Neolamprologus brichardi            (69) KYRDMRTTTNLYLCSMAVSDLLIFLCMPLDLYRMWRYRPWRFGDALCKLFQFVSESSTYSTILSITALSVERYLAICFPLRAKALVTKRRVRALICLLWT
```

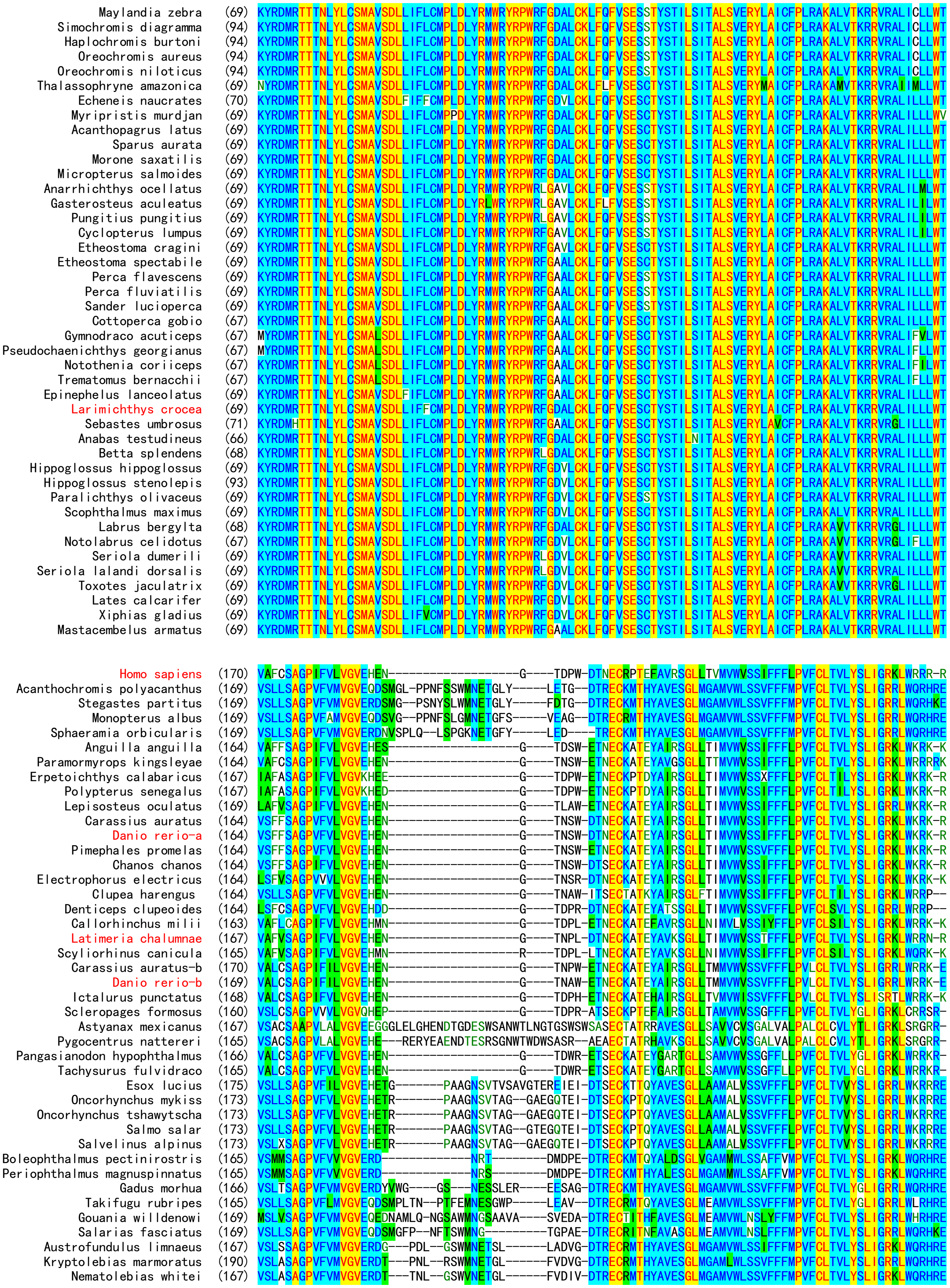
Maylandia zebra (69)
Simochromis diagramma (94)
Haplochromis burtoni (94)
Oreochromis aureus (94)
Oreochromis niloticus (94)
Thalassophryne amazonica (69)
Echeneis naucrates (70)
Myripristis murdjan (69)
Acanthopagrus latus (69)
Sparus aurata (69)
Morone saxatilis (69)
Micropterus salmoides (69)
Anarrhichthys ocellatus (69)
Gasterosteus aculeatus (69)
Pungitius pungitius (69)
Cyclopterus lumpus (69)
Etheostoma cragini (69)
Etheostoma spectabile (69)
Perca flavescens (69)
Perca fluviatilis (69)
Sander lucioperca (69)
Cottoperca gobio (67)
Gymnodraco acuticeps (67)
Pseudochaenichthys georgianus (67)
Notothenia coriiceps (67)
Trematomus bernacchii (67)
Epinephelus lanceolatus (69)
Larimichthys crocea (69)
Sebastes umbrosus (71)
Anabas testudineus (66)
Betta splendens (68)
Hippoglossus hippoglossus (69)
Hippoglossus stenolepis (93)
Paralichthys olivaceus (69)
Scophthalmus maximus (69)
Labrus bergylta (68)
Notolabrus celidotus (67)
Seriola dumerili (69)
Seriola lalandi dorsalis (69)
Toxotes jaculatrix (69)
Lates calcarifer (69)
Xiphias gladius (69)
Mastacembelus armatus (69)
Homo sapiens (170)
Acanthochromis polyacanthus (169)
Stegastes partitus (169)
Monopterus albus (169)
Sphaeramia orbicularis (169)
Anguilla anguilla (164)
Paramormyrops kingsleyae (164)
Erpetoichthys calabaricus (167)
Polypterus senegalus (167)
Lepisosteus oculatus (169)
Carassius auratus (164)
Danio rerio-a (164)
Pimephales promelas (164)
Chanos chanos (164)
Electrophorus electricus (164)
Clupea harengus (164)
Denticeps clupeoides (164)
Callorhinchus milii (163)
Latimeria chalumnae (167)
Scyliorhinus canicula (165)
Carassius auratus-b (170)
Danio rerio-b (169)
Ictalurus punctatus (168)
Scleropages formosus (160)
Astyanax mexicanus (167)
Pygocentrus nattereri (165)
Pangasianodon hypophthalmus (166)
Tachysurus fulvidraco (165)
Esox lucius (175)
Oncorhynchus mykiss (173)
Oncorhynchus tshawytscha (173)
Salmo salar (173)
Salvelinus alpinus (173)
Boleophthalmus pectinirostris (165)
Periophthalmus magnuspinnatus (165)
Gadus morhua (166)
Takifugu rubripes (165)
Gouania willdenowi (169)
Salarias fasciatus (169)
Austrofundulus limnaeus (167)
Kryptolebias marmoratus (190)
Nematolebias whitei (167)

```
Cyprinodon tularosa             (168) VSLFSAGPVFVLVGVERDSNEQSN-------------------TLDVE-DTRECKMTQYAVESGLMGAMVWLSSVFFFMPVFCLTVLYSLIGRRLWQRHRE
Cyprinodon variegatus           (168) VSLFSAGPVFVLVGVERDSNEQSN-------------------TLDVE-DTRECRMTQYAVESGLMGAMVWLSSVFFFMPVFCLTVLYSLIGRRLWQRHRE
Fundulus heteroclitus           (169) VSLCSAGPVFALVGVEMDTMEQNLE--SNET------------VFDVG-DTRECRMTHYAVESGLMGAMVWLSSVFFFMPVFCLTVLYSLIGRRLWQRHRE
Poecilia formosa                (169) VSLFSAGPVFVMVGVERDSVEPLNL--TSRINATGS-------LLDVG-DTRECRMTHYAVESGLMGAMVWLSSVFFFMPVFCLTVLYSLIGRRLWQRHRE
Poecilia mexicana               (169) VSMFSAGPVFVMVGVERDSVEPLNL--TSRINATGS-------LLDVG-DTRECRMTHYAVESGLMGAMVWLSSVFFFMPVFCLTVLYSLIGRRLWQRHRE
Poecilia latipinna              (169) VSLFSAGPVFIMVGVERDSVEPLNL--TSRINATGS-------LLDVG-DTRECRMTHYAVESGLMGAMVWLSSVFFFMPVFCLTVLYSLIGRRLWQRHRE
Poecilia reticulata             (169) VSLFSAGPVFVMVGVERDTVEPLNL--TSRINTTGL-------LLDVG-DTRECRMTHYAVESGLMGAMVWLSSVFFFMPVFCLTVLYSLIGRRLWQRHRE
Xiphophorus couchianus          (169) VSLFSAGPVFVMVGVERDNVEPLNL--TSRINTTGL-------LLDVG-DTRECRMTHYAVESGLMGAMVWLSSVFFFMPVFCLTVLYSLIGRRLWQRHRE
Xiphophorus hellerii            (169) VSLFSAGPVFVMVGVERDNVEPLNL--TSRINTTGL-------LLDVG-DTRECRMTHYAVESGLIGAMVWLSSVFFFMPVFCLTVLYSLIGRRLWQRHRE
Xiphophorus maculatus           (169) VSLFSAGPVFVMVGVERDNVEPLNL--TSRINTTGL-------LLDVG-DTRECRMTHYAVESGLMGAMVWLSSVFFFMPVFCLTVLYSLIGRRLWQRHRE
Nothobranchius furzeri          (169) VSLMSAGPVFAMVGVERDYVDPTNL--R-WMNETGL-------FLDVG-DTRECRMTHYAVQSGLMEAMVWLSSVFFFMPVFCLTMLYSLIGRRLWQRQRE
Oryzias latipes                 (194) VSLLSAGPVFVMVGVERDITEPPNF--SSWINETGL-------FLDVG-DTRECRMTHYAVESGLMEAMVWLSSVFFFMPVFCLTVLYSLIGRRLWQRHRE
Oryzias melastigma              (195) VSLLSAGPVFVMVGVERDITEPPNF--SSWINGTGL-------FLDVG-DTRECRMTHYAVESGLMEAMVWLSSVFFFMPVFCLTVLYSLIGRRLWQRHRE
Cynoglossus semilaevis          (172) VSLLSAGPVFVMVGVERDGTGMEPLNFTLDVNETDLY------PEIG--DTRECKMTHYAVESGLMEAMVWLSSVFFFMPVFCLTVLYSLIGRRLWQRHRE
Hippocampus comes               (170) VAVLSAGPVFVMVGVERDYPG-----FD--ANDTLRMD--EEEEEYG-DTRECKMTRYAVESGLMGAMVWLSSVFFFMPVFCLTVLYSLIGRRLWRRHRE
Syngnathus acus                 (170) VSLLSAGPVFVMVGVERDHLS-----FS--ANDTHALG--REEDEYR-DTRECKMTHYAVESGLMGAMVWLSSVFFFMPVFCLTVLYSLIGRRLWHRHRN
Parambassis ranga               (169) VSLLSAGPVFVMVGVERDNIGPN---FSLWMNESGLN------FEIG--DTRECKMTQYAVESGLMGAMVWLSSVFFFMPVFCLTVLYSLIGRRLWQRHRE
Archocentrus centrarchus        (197) VSLLSAGPVFVMVGVEQDTMGSL--NFSSWMNETSLF------LEAE--DTRECKMTHYAVQSGLMEAMVWLSSVFFFMPVFCLTVLYSLIGRRLWQRHRE
Astatotilapia calliptera        (194) VSLLSAGPVFVMVGVEQDTMGPL--NFSSWMNESNLF------LEAE--DTRECKMTHYAVQSGLMGAMVWLSSVFFFMPVFCLTVLYGLIGRRLWQRHRE
Pundamilia nyererei             (194) VSLLSAGPVFVMVGVEQDTMGPL--NFSSWMNESNLF------LEAE--DTRECKMTHYAVQSGLMGAMVWLSSVFFFMPVFCLTVLYGLIGRRLWQRHRE
Neolamprologus brichardi        (169) VSLLSAGPVFVMVGVEQDTMGPL--NFSSWMNESNLF------LEAE--DTRECKMTHYAVQSGLMGAMVWLSSVFFFMPVFCLTVLYGLIGRRLWQRHRE
Maylandia zebra                 (169) VSLLSAGPVFVMVGVEQDTMGPL--NFSSWMNESNLF------LEAE--DTRECKMTHYAVQSGLMGAMVWLSSVFFFMPVFCLTVLYGLIGRRLWQRHRE
Simochromis diagramma           (194) VSLLSAGPVFVMVGVEQDTMGPL--NFSSWVNESNLF------LEAE--DTRECKMTHYAVQSGLMGAMVWLSSVFFFMPVFCLTVLYGLIGRRLWQRHRE
Haplochromis burtoni            (194) VSLLSAGPVFVMVGVEQDTMGPL--NFSSWMNESNLF------LEAE--DTRECKMTHYAVQSGLMGAMVWLSSVFFFMPVFCLTVLYGLIGRRLWQRHRE
Oreochromis aureus              (194) VSLLSAGPVFVMVGVEQDTMGPL--NFSSWMNETNLF------LETE--DTRECKMTHYAVQSGLMGAMVWLSSVFFFMPVFCLTVLYSLIGRRLWQRHRE
Oreochromis niloticus           (194) VSLLSAGPVFVMVGVEQDTMGPL--NFSSWMNETNLF------LETE--DTRECKMTHYAVQSGLMGAMVWLSSVFFFMPVFCLTVLYSLIGRRLWQRHRE
Thalassophryne amazonica        (169) VSLLSAGPVFVMVGVEQDSVG----HPSAEMNENGFV------LEAG--DTRECKMTHYALESGLMGAMVWLSSVFFFMPVFCLTVLYSLISRRLWQRHRE
Echeneis naucrates              (170) VSLLSAGPVFVMVGVERDSMAPQ--NISSGMNGTGFP------LEDG--DTRECKMTHYAVESGLIGAMVWLSSVFFFMPVFCLTVLYSLIGRRLWQRHRE
Myripristis murdjan             (169) VSLLSAGPVFVMVGVERD--------NSSWGVNESGYS-----LEVR--DTRECKMTHYAVESGLMGAMVWLSSVFFFMPVFCLTVLYSLIGRRLWQRHRE
Acanthopagrus latus             (169) VSLLSAGPVFVMVGVERDSMWPGN-LSWVGMNGTGFF------PEEG--DTRECKMTHYAVESGLMGAMVWLSSVFFFMPVFCLTVLYSLIGRRLWQRHRE
Sparus aurata                   (169) VSLLSAGPVFVMVGVERDSMWQGN-LSWV-VNGTGFF------PEEG--DTRECKMTHYAVESGLMGAMVWLSSVFFFMPVFCLTVLYSLIGRRLWQRHRE
Morone saxatilis                (169) VSLLSAGPVFVMVGVERDSMWPTN-LSSG-MNETGFY------LEAG--DTRECKMTHYAVESGLMGAMVWLSSVFFFMPVFCLTVLYSLIGRRLWQRHRE
Micropterus salmoides           (169) VSLLSAGPVFVMVGVERDNMGPTN-FSSG-MNETGFF------LEAG--DTRECKMTHYAVESGLMGAMVWLSSVFFFMPVFCLTVLYSLIGRRLWQRHRE
Anarrhichthys ocellatus         (169) VSLFSAGPVFVMVGVERDSMWPN--FSLGINETDFS-------LED----TRECKMTHYAVESGLMGAMVWLNSVFFFIPVFCLTVVYSLIGRRLWQRNRE
Gasterosteus aculeatus          (169) VALFSAGPVFIMVGVERDSVWSN---LGSGMNYTDFS------LEN----TRECKITHYAVVSGLVEAMVWLNSVFFFIPVFCLTVVYSLIGRRLWQRNRE
Pungitius pungitius             (169) VALVSAGPVFIMVGVERDSVWPN---LSSGMNDSDS--------------TRECKFTHFAVESGLMGAMVWLNSIFFFVPVFCLTVVYSLIGRRLWQRNRE
Cyclopterus lumpus              (169) VSLVSAGPVFIMVGVERDGASPGG--VISAMNETDFS------PEAG--DTRECTMTHYAVESGLMGAMVWLNSVFFFMPVFCLTVLYSLIGRRLWQRDRE
Etheostoma cragini              (169) VSLLSAGPVFVMVGVERDNMGPH---FSSGMNYTGFF------LEAG--DTRECKMTHYAVESGLMGAMVWLSSVFFFMPVFCLTVLYSLIGRRLWQRHRE
Etheostoma spectabile           (169) VSLLSAGPVFVMVGVERDNMGPH---FNLGMNDTDFS------LEAG--DTRECKMTHYAVESGLIGAMVWLSSVFFFMPVFCLTVLYSLIGRRLWQRHRE
Perca flavescens                (169) VSLLSAGPVFVMVGVERDSMGPH---FSSEMNDTDFS------LEAG--DTRECKMTHYAVESGLMGAMVWLSSVFFFMPVFCLTVLYSLIGRRLWQRHRE
Perca fluviatilis               (169) VSLLSAGPVFVMVGVESDSMGPH---FSSEINDTDFS------LEAG--DTRECKMTHYAVESGLMGAMVWLSSVFFFMPVFCLTVLYSLIGRRLWQRHRE
Sander lucioperca               (169) VSLMSAGPVFVMVGVERDSMGPH---FSSGMNDTDFS------LEAG--DTRECKMTHYAVESGLMGAMVWLSSVFFFMPVFCLTVLYSLIGRRLWQRHRE
Cottoperca gobio                (167) VSLMSAGPVFVMVGVECDSMWPN---FSAAMNETGFS------PEAG--DTRECKITHYAVESGLMGAMVWLSSVFFFMPVFCLTVLYSLIGRRLWQRHRE
Gymnodraco acuticeps            (167) VSVMSAGPVFVMVGVERD-MSPH---FSSEMNETGYI------LEPG--DTRECKITHFAVESGLMGAMVWLSSVFFFMPVFCLTVLYSLIGRRLWQRHRE
Pseudochaenichthys georgianus   (167) VSVMSAGPVFVMVGVERD-MSPH---FSPEMNETGYV------LEPG--DTRECKITHFAVESGLMGAMVWLSSVFFFMPVFCLTVLYSLIGRRLWQRHRE
Notothenia coriiceps            (167) VSVMSAGPVFVMVGVERDSMSPH---FSPEMNETGYF------LEPG--DTRECKITHFAVESGLMGAMVWLSSVFFFIPVFCLTVLYSLIGRRLWQRHRE
Trematomus bernacchii           (167) VSVMSAGPVFVMVGVERDSMSPN---VSLEMNETGYF------VEPG--DTRECKITHFAVESGLMGAMVWLSSVFFFMPVFCLTVLYSLIGRRLWQRHRE
Epinephelus lanceolatus         (169) VSLLSAGPVFVMVGVERDSMWPD---YSSGMNETGFS------PEET--DTRECKMTHYAVESGLMGAMVWLSSVFFFMPVFCLTVLYSLIGRRLWQRHRE
Larimichthys crocea             (169) VSLLSAGPVFVMVGVERDSMWPTN-FTSE-MNETDSA------LED----TRECKMTHYAVESGLMGAMVWLSSVFFFMPVFCLTVLYSLIGRRLWQRHRE
Sebastes umbrosus               (171) VSLLSAGPVFVMVGVERDSMGPN---FTWEMNETGLF------MED----TRECKMTHYAVESGLMGAMVWLSSVFFFMPVFCLTVLYSLIGRRLWQRHRE
Anabas testudineus              (166) VSLLSAGPVFVMVGVERDSMGPTN--FTLGMNETEFF------LEVG--DTRECKMTHYAVESGLMGAMVWLSSVFFFMPVFCLTVLYSLIGRRLWQRHRE
Betta splendens                 (168) VSLLSAGPVFVMVGVERDSVGATN--FSSGTNETELV------PEVG--DTRECKMTHYAVESGLMGAMVWLSSVFFFMPVFCLTVLYSLIGRRLWQRHRE
Hippoglossus hippoglossus       (169) VSLLSAGPVFVMVGVERDIMGP----FGSDMNETGFS------VEGGG-DTRECRMTHYAVESGLMGAMVWLSSVFFFMPVFCLTVLYSLIGRRLWQRHRE
Hippoglossus stenolepis         (193) VSLLSAGPVFVMVGVERDIMGP----FGSDMNETGFS------VEGGG-DTRECRMTHYAVESGLMGAMVWLSSVFFFMPVFCLTVLYSLIGRRLWQRHRE
Paralichthys olivaceus          (169) VSLLSAGPVFVMVGVERDSMGP----FSSDMNETGLS------LEGGG-DTRECKMTHYAVESGLMGAMVWLSSVFFFMPVFCLTVLYSLIGRRLWQRHRE
Scophthalmus maximus            (169) VSLLSAGPVFVMVGVERDSVGPPY--FSSGANETDFF------AEDG--DTRECKMTHYAVESGLMGAMVWLSSVFFFMPVFCLTVLYSLIGRRLWQRHRE
Labrus bergylta                 (168) VSLLSAGPVFVMVGVERDSSG----------MNETVYS-----LEEV--DTRECKMTHYAVESGLMGAMVWLSSVFFFMPVFCLTVLYSLIGRRLWQRHRE
Notolabrus celidotus            (167) VSLVSAGPVFVMVGVERDIMGPPN--YSSGLNETDFY------LEDG--DTRECKMTHYAVESGLMGAMVWLSSVFFFMPVFCLTVLYSLIGRRLWQRHRE
Seriola dumerili                (169) VSLLSAGPVFVMVGVERDSMGPLN--LSSGKNETGSS------LEGE--DTRECKMTHYAVESGLMGAMVWLSSVFFFMPVFCLTVLYSLIGRRLWQRHRE
Seriola lalandi dorsalis        (169) VSLLSAGPVFVMVGVERDSMGPLN--MSSGKNETGSS------LEGE--DTRECKMTHYAVESGLMGAMVWLSSVFFFMPVFCLTVLYSLIGRRLWQRHRE
Toxotes jaculatrix              (169) VSLLSAGPVFVMVGVERDSMGPPN--FSSGMNETGSL------LEGG--DTRECKMTHYAVESGLMGAMVWLSSVFFFMPVFCLTVLYSLIGRRLWQRHRE
Lates calcarifer                (169) VSLLSAGPVFVMVGVERDYMGPPN--FSSGMNETGFF------QDGG--DTRECKMTHYAVESGLMGAMVWLSSVFFFMPVFCLTVLYSLIGRRLWQRHRE
Xiphias gladius                 (169) VSLLSAGPVFVMVGVERDSMVPPN--FSSGMNETDFF------LEGG--DTRECKMTHYAVVSGLMGAMVWLSSVFFIMPVFCLTVLYTLIGRRLWQRHRE
Mastacembelus armatus           (169) VSLLSAGPVFVMVGVEQDSMGPPN--FSLGMNETGFS------LVVG--DTRECKMTHYAVESGLMGAMVWLSSVFFFMPVFCLTVLYSLIGRRLWQRHRE

                                                                           6.58
Homo sapiens                    (245) GDAVVGASLRDQNHKQTVKMLAVVVFAFILCWLPFHVGRYLFSKSFEP-GSLEIAQISQYCNLVSFVLFYLSAAINPILYNIMSKKYRVAVFRLLGFEP-
Acanthochromis polyacanthus     (261) TNMSSRVAHRDKSNRQTIKMLVVVVLAFVLCWLPFHVGRYLQFRSLDA-PSPLLSALSEYCSLVSVVLFYLSAAINPILYNTMSWKYRGAAARLFGLTD-
Stegastes partitus              (260) TNMSSRVAHRDKSNRQTIKMLVVVVLAFVLCWLPFHVGRYLQFRSLDA-PSPLLSLLSEYCSLVSVVLFYLSAAINPILYNTMSWKYRGAAARLFGLTD-
Monopterus albus                (260) TNISSRVAHRDKSNRQTIKMLVVVVLAFVLCWLPFHVGRYLQFRSLDA-PSPLLSLLSEYCSLVSVVLFYLSAAINPILYNTMSWKYRGAAARLFGLTD-
Sphaeramia orbicularis          (258) TNMSSRVAHRDKSNRQTIKMLVVVVLAFVLCWLPFHVGRYLQFRSLDA-PSRLLSTLSEYCSLVSVVLFYLSAAINPILYNTMSWKYRGAAARLFGLTD-
Anguilla anguilla               (239) ETIGPNISSRDKNNKQTVKMLAVVVFAFVLCWLPFHVGRYLISKSSEA-GSPLMSLISEYCNLVSFVLFYLSAAINPILYNIMSKKYRIAACRLFGLKQ-
Paramormyrops kingsleyae        (240) EKIGPNISSRDKNNKQTVKMLAVVVFAFVLCWLPFHVGRYLISKSSEA-GSPLVSLISQYCNLVSFVLFYLSAAINPILYNIMSKKYRDAACKLFRGKH-
Erpetoichthys calabaricus       (242) DTIGPNMSSRDKNNKQTVKMLAVVVFAFVLCWLPFHVGRYLFSKSSEA-NSLEMSQMSQYCNLVSFVLFYLSAAINPILYNIMSKKYRVAACKLFGLKQ-
Polypterus senegalus            (242) DTIGPNMSSRDKNNKQTVKMLAVVVFAFVLCWLPFHVGRYLFSKSSEA-NSLEMSQMSQYCNLVSFVLFYLSAAINPILYNIMSKKYRVAACKLFGLKQ-
Lepisosteus oculatus            (244) EAIGPNISSRDKNNKQTVKMLAVVVFAFVLCWLPFHVGRYLFSKSSEA-GSPLMSQISQYCNLVSFVLFYLSAAINPILYNIMSKKYRVAACKLFGLKQ-
Carassius auratus               (239) ETIGENASSRDKNNRQTVKMLAVVVFAFVLCWLPFHVGRYLISKSTEM-GSPVMSVISHYCNLISFVLFYLSAAINPILYNIMSKKYRMAACKLFGLHH-
Danio rerio-a                   (239) ETIGENASSRDKSNRQTVKMLAVVVFAFVLCWLPFHVGRYLISKSTEM-GSPVMSIISHYCNLISFVLFYLSAAINPILYNIMSKKYRMAACKLFGLRN-
Pimephales promelas             (239) ETIGENASSRDKSNRQTVKMLAVVVFAFVLCWLPFHVGRYLISKSTEM-GSPVMSVISQYCNLISFVLFYLSAAINPILYNIMSKKYRMAACKLFGLRN-
Chanos chanos                   (239) ETIGPNISNREKNNRQTVKMLAVVVFAFVLCWLPFHVGRYLISKSSEV-GSPVLSLISQYCNLISFVLFYLSAAINPILYNIMSKKYRSAACKLFGLNR-
Electrophorus electricus        (239) ETAGTNLSSRDKNNRQTVKMLAVVVFAFVLCWLPFHVGRYLISKSSQA-VSPIMSLITQYCSIVSFVLFYLSAAINPILYNIMSKKYRIAACRLLSLRR-
Clupea harengus                 (238) -----NISSRDKNNKQTVKMLVVVVFAFVLCWLPFHVVRYLISKSSET-GSPMMADITQYCSLISFVLFYLSAAINPILYNIMSKKYRVAACTLFGVKK-
Denticeps clupeoides            (238) -----GASSRDKNNRQTVKMLVVVVFAFVLCWLPFHVGRYLVSKTSES-GSPLMSTISQYCNLISFVLFYLSAAINPILYNIMSKKYRVAACKLLGLNR-
Callorhinchus milii             (238) ETIGPNLNNRHKNNKQTIKMLAVVVFAFILCWLPFHVGRYLFSNSE------AGSQISQYCNLVSLVLFYLSAAINPILYNIMSKKFRVAAFKMFRVKK-
Latimeria chalumnae             (242) ETIGPNICIRDRNNKQTVKMLAVVVFAFVLCWLPFHVGRYLFSKSTEV-GSFEMSQISQYCNLVSFVLFYLSAAINPILYNIMSKKYRVAACKLFGVQQ-
Scyliorhinus canicula           (240) ENIRPSVSSREKNNKQTVKMLAVVVFAFVLCWLPFHLGRYLFTKSSEI-GSYEMLQISQYCSLVSFVLFYLSAAINPILYNIMSKKYRVAALKLFHVKQ-
```

```
Carassius auratus-b            (245) NPVGPISS-REKNNKQTVKMLAVVVLAFVLCWLPFHVGRYLFSKSSEA-NSPLISQISEYCNLVSFVLFYLSAAINPILYNIMSKKYRSAACKLFGVKR-
Danio rerio-b                  (244) NPVGPISS-RDKSNKQTVKMLAVVVLAFVLCWLPFHVGRYLVSKSSEA-NSPVISQISEYCNLVSFVLFYLSAAINPILYNIMSKKFRSAACKLFRVKR-
Ictalurus punctatus            (243) NPVGPVSS-REKNNTQTVKMLAVVVFAFVLCWLPFHLGRYLFSKSSEA-DSPLITQMSEYCNLVSLVLFYFSAAINPILYNIMSKKYRIMACRLFGVRC-
Scleropages formosus           (235) ATPGAHASVRDKSNRQTVRMLAVVVFAFVLCWLPFHVGRYLISKSSE-AVSPLLSQITQYCNLVSFVLFYLSAAINPVLYNIMSRKYRTAACQLFGVKR-
Astyanax mexicanus             (266) --ADARSSAREKSHRQSVKLLAVVVLAFVLCWIPFHVVRFLISTSSAA-GSPIMSLISQYCSLISFVLFYLSAAINPILYNIMSEKYRAAVCRLFGIQG-
Pygocentrus nattereri          (259) --ADARSSARDKNHRQSIKLLAVVVLAFVLCWLPFHVVRFLISASSAA-GSPIMSLISQYCSLISFVLFYLSAAINPILYNIMSEKYRAAVCRLFGIHS-
Pangasianodon hypophthalmus    (241) -RRRDRMKSRDSSNRQTIKMLAVMVFAFVLCWLPFHVGRYLFSASPEAFASPLWSLIAQYCSLVSFVLFYLSAAINPILYNAMSKKYRNATLRLFSHSS-
Tachysurus fulvidraco          (240) -RRRDRMKSRDSSNRQTIKMLAVMVFAFVLCWLPFHVGRYLFSASPEAFVSPLWSLVSQYCSLVSFVLFYLSAAINPILYNTMSKKYRDATLRLFSHSS-
Esox lucius                    (267) NNIGANVAHRDKSNRQTVKMLAVVVFAFVLCWLPFHLQRYLMSHSSEG-SSPIWSLITQYCSLVSTVLFYLSAAINPVLYNTMSRKYRSAAAQLFGLQE-
Oncorhynchus mykiss            (263) NNIGANVAHRDKSNRQTVKMLAVVVFAFVLCWLPFHLHRYLMSHSSEG-SSPLWSLFTQYCSLVSTVLFYLSAAINPVLYNTMSRKYRSAAAQLFGLQE-
Oncorhynchus tshawytscha       (263) NNIGANVAHRDKSNRQTVKMLAVVVFAFVLCWLPFHLHRYLMSHSSEG-SSPLWSLFTQYCSLVSTVLFYLSAAINPVLYNTMSRKYRSAAAQLFGLQE-
Salmo salar                    (263) NNIGANVAHRDKSNRQTVKMLAVVVFAFVLCWLPFHLHRYLMSHSSEG-SSPLWSLFTQYCSLVSTVLFYLSAAINPVLYNTMSRKYRSAAAQLFGLQE-
Salvelinus alpinus             (263) NNIGANVAHRXKSNRQTVKMLVVGSLPSVLCWLPFHLHRYLMSHSSEG-SSPLWSLFTQYCSLVSTVLFYLSAAINPVLYNTMSRKYRSAAAQLFGLQE-
Boleophthalmus pectinirostris  (243) TCMSSRIAHRERSNRQTIKMLVVVVFAFVLCWLPFHVGRYLQFRSLNT-QSQLLSTLSGYCSLVSVVLFYLSAAINPILYNTMSWKYRRAAARLFGLTDN
Periophthalmus magnuspinnatus  (243) TCMSSRVAHRERSNRQTIKMLVVVVFAFVLCWLPFHVGRYLQFRSLYT-QSQLLSTLSGYCSLVSVVLFYLSAAINPILYNTMSWKYRRAAGRLFGITDN
Gadus morhua                   (254) NIMGSRVAHRDKSNRQTIKMLVVVVFAFVLCWLPFHVGRYLQFRSLDA-PSPMLSALSEYCSLVSVVLFYLSAAVNPILYNTMSWKYRCAAVRLFGLTE-
Takifugu rubripes              (256) TTINSRVAYRDKSNRQTIKMLVVVVLAFVLCWLPFHVGRYLQFRSLDA-PSPLLSLLSEYCSLVSVVLFYLSAAINPILYNTMSWKYRSAAARLFGVSE-
Gouania willdenowi             (263) TNMSTRMAHRDKSNRQTIKMLVVVVLAFILCWLPFHVGRYLQFRYLDA-PSQLLMALSQYCGMVSVVLFYLSAAINPILYNTMSWKYRGAAARLFGLAD-
Salarias fasciatus             (257) TNISSRVAHRDKSNRQTIKMLVVVVLAFVLCWLPFHVGRYLHFRFLDS-PSPLLLVLSQYFSMVSVVLFYLSAAINPILYNTMSWKYRSAAARLFGLAD-
Austrofundulus limnaeus        (255) TNMSSRVSYRDKSNRQTIKMLVVVVLAFVLCWLPFHVGRYLQFRSLDA-PSPLLSLLSEYCSLLSVVLFYLSAAINPILYNTMSWKYRSAAARLFGLAD-
Kryptolebias marmoratus        (278) TNMSSRVSHRDKSNRQTIKMLVVVVLAFVLCWLPFHVGRYLQFRSLDA-PSPLLSVLSEYCSLLSVVLFYLSAAINPILYNTMSWKYRGAAARLFGLVD-
Nematolebias whitei            (255) TNMSSRVSHRDKSNRQTIKMLVVVVLAFVLCWLPFHVGRYLQFRSLDA-PSPLLSVLSEYCSLLSVVLFYLSAAINPILYNTMSWKFRRAAARVFGLED-
Cyprinodon tularosa            (249) TNMSSRVAHRDKSNRQTIKMLVVVVLAFVLCWLPFHVGRYLQFRSLDA-PSPLLSVLSEYCSLLSVVLFYLSAAINPILYNTMSWKYRGAAARLFGLTD-
Cyprinodon variegatus          (249) TNMSSRVAHRDKSNRQTIKMLVVVVLAFVLCWLPFHVGRYLQFRSLDA-PSPLLSVLSEYCSLLSVVLFYLSAAINPILYNTMSWKYRGAAARLFGLTD-
Fundulus heteroclitus          (255) TSMSSRVAHRDKSNRQTIKMLVVVVLAFVLCWLPFHVGRYLQFRSLDA-PSPLLSVLSEYCSLLSVVLFYLSAAINPILYNTMSWKYRGAAARLFGLTD-
Poecilia formosa               (260) TNMSSRVAHRDKSNRQTIKMLVVVVLAFVLCWLPFHVGRYLQFRSLDA-PSPLLSVLSEYCSLLSVVLFYLSAAVNPILYNTMSWKYRGAAARLFGLTD-
Poecilia mexicana              (260) TNMSSRVAHRDKSNRQTIKMLVVVVLAFVLCWLPFHVGRYLQFRSLDA-PSPLLSVLSEYCSLLSVVLFYLSAAVNPILYNTMSWKYRGAAARLFGLTD-
Poecilia latipinna             (260) TNMSSRVAHRDKSNRQTIKMLVVVVLAFVLCWLPFHVGRYLQFRSLDA-PSPLLSVLSEYCSLLSVVLFYLSAAVNPILYNTMSWKYRGAAARLFGLTD-
Poecilia reticulata            (260) TNMSSRVAHRDKSNRQTIKMLVVVVLSFVLCWLPFHVGRYLQFRSLDA-PSPLLSVLSEYCSLLSVVLFYLSAAINPILYNTMSWKYRVAAARLFGLTD-
Xiphophorus couchianus         (260) TNMSSRVAHRDKSNRQTIKMLVVVVLAFVLCWLPFHVGRYLQFRFLDA-PSPLLSVLSEYCSLLSVVLFYLSAAINPILYNTMSWKYRGAAARLFGLTD-
Xiphophorus hellerii           (260) TNMSSRVAHRDKSNRQTIKMLVVVVLAFVLCWLPFHVGRYLQFRFLDA-PSPLLSLLSEYCSLLSVVLFYLSAAINPILYNTMSWKYRGAAARLFGLTD-
Xiphophorus maculatus          (260) TNMSSRVAHRDKSNRQTIKMLVVVVLAFVLCWLPFHVGRYLQFRFLDA-PSPLLSLLSEYCSLLSVVLFYLSAAINPILYNTMSWKYRGAAARLFGLTD-
Nothobranchius furzeri         (259) TNVSSRVSHRDKSNRQTIKMLVVVVLAFVLCWLPFHVGRYLQFRSLDA-PSPLLSVLSEYCSLVSVVLFYLSAAINPILYNTMSWKYRGAAARLFGLAD-
Oryzias latipes                (285) TSVSSRVAHRDKSNRQTIKMLVVVVLAFVLCWLPFHVGRYLQFRSLDA-PSPLLSVLSEYCSMVSVVLFYLSAAINPVLYNTMSWKYRGAAARLFGLAD-
Oryzias melastigma             (286) TSVSSRVAHRDKSNRQTIKMLVVVVLAFVLCWLPFHVGRYLQFRSLDA-PSPLLSVLSEYCSLVSVVLFYLSAAINPVLYNTMSWKYRGAAARLFGLAD-
Cynoglossus semilaevis         (265) TNIRSRVSYRDKSNRQTIKMLVVVVLAFVLCWLPFHVGRYLQFRSLDA-PSPFLSLLSEYCSLVSVVLFYLSAAINPILYNTMSWKYRVAAARLFGLAD-
Hippocampus comes              (260) TNISSCVAHRDKCNRQTVKMLVVVVLAFVLCWLPFHVCRYLQFRSLDA-PSPLLSALSEHCSLLSVVLFYLSAAINPILYNTMSWKYRGAAARLFGLAG-
Syngnathus acus                (260) TNIR---SHRDRSNRQTIKMLVVVVLAFVLCWLPFHVCRYLQSRSLDA-PSPLLSALSEYCSLLSVVLFYLSAAINPILYNTMSWKFRGAAARLFGLGH-
Parambassis ranga              (259) TNMTSRVVHRDKSNRQTIKMLVVVVLAFVLCWLPFHVGRYLQFRSLDA-PSPLLSVLSEYCSLVSVVLFYLSAAINPILYNTMSWKYRSAAARLFGLTD-
Archocentrus centrarchus       (288) TNMSSRVSHRDKSNRQTIKMLVVVVLAFVLCWLPFHVGRYLQFRSLDA-PSPLLSVLSEYCSLVSVVLFYLSAAINPILYNTMSWKYRGAAARLFGLTY-
Astatotilapia calliptera       (285) TNMSNRVSHRDKSNRQTIKMLVVVVLAFVLCWLPFHVGRYLQFRSLDA-PSPLLSLLSEYCSLVSVVLFYLSAAINPILYNTMSWKYRGAAARLFGLTD-
Pundamilia nyererei            (285) TNMSNRVSHRDKSNRQTIKMLVVVVLAFVLCWLPFHVGRYLQFRSLDA-PSPLLSLLSEYCSLVSVVLFYLSAAINPILYNTMSWKYRGAAARLFGLTD-
Neolamprologus brichardi       (260) TNMSNRVSHRDKSNRQTIKMLVVVVLAFVLCWLPFHVGRYLQFRSLDA-PSPLLSLLSEYCSLVSVVLFYLSAAINPILYNTMSWKYRGAAARLFGLTD-
Maylandia zebra                (260) TNMSNRVSHRDKSNRQTIKMLVVVVLAFVLCWLPFHVGRYLQFRSLDA-PSPLLSLLSEYCSLVSVVLFYLSAAINPILYNTMSWKYRGAAARLFGLTD-
Simochromis diagramma          (285) TNMSNRVSHRDKSNRQTIKMLVVVVLAFVLCWLPFHVGRYLQFRSLDA-PSPLLSLLSEYCSLVSVVLFYLSAAINPILYNTMSWKYRGAAARLFGLTD-
Haplochromis burtoni           (285) TNMSNRVSHRDKSNRQTIKMLVVVVLAFVLCWLPFHVGRYLQFRSLDA-PSPLLSLLSEYCSLVSVVLFYLSAAINPILYNTMSWKYRGAAARLFGLTD-
Oreochromis aureus             (285) TNMSNRVSHRDKSNRQTIKMLVVVVLAFVLCWLPFHVGRYLQFRSLDA-PSPLLSLLSEYCSLVSVVLFYLSAAINPILYNTMSWKYRGAAARLFGMTD-
Oreochromis niloticus          (285) TNMSNRVSHRDKSNRQTIKMLVVVVLAFVLCWLPFHVGRYLQFRSLDA-PSPLLSLLSEYCSLVSVVLFYLSAAINPILYNTMSWKYRGAAARLFGLTD-
Thalassophryne amazonica       (258) TNVNSRVAHRDKSNRQTIKMLVVVVLAFVLCWLPFHVGRYLQFRSLDA-PSPLLSLLSEYCTLVSVVLFYLSAAINPILYNTMSWKYRGAAARLFGLTD-
Echeneis naucrates             (261) TNSSSRVAHRDKSNRQTIKMLVVVVLAFVLCWLPFHVGRYLQFRSLDA-PSPLLSVLSEYCSLVSVVLFYLSAAINPILYNTMSWKYRGAAARLFGLTD-
Myripristis murdjan            (255) TNIGSRVAHRDKSNRQTIKMLVVVVLAFVLCWLPFHVGRYLQFRSMDA-PSPLLSVLSEYCTLVSVVLFYLSAAINPILYNTMSWKYRGAAARLFGLTD-
Acanthopagrus latus            (261) TNINSRVAHREKSNRQTIKMLVVVVLAFVLCWLPFHVGRYLQFRSLDA-PSPLLSLLSEYCSLVSVVLFYLSAAINPILYNIMSWKYRGAAARLFGLID-
Sparus aurata                  (260) TNINSRVAHREKSNRQTIKMLVVVVLAFVLCWLPFHVGRYLQFRSLDA-PSPLLSLLSEYCSLVSVVLFYLSAAINPILYNIMSWKYRGAAARLFGLTD-
Morone saxatilis               (260) TNINSRVAHRDKSNRQTIKMLVVVVLAFVLCWLPFHVGRYLQFRSLDA-PSPLLSVLSEYCSLVSVVLFYLSAAINPILYNTMSWKYRGAAARLFGLTD-
Micropterus salmoides          (260) TNISSRVAHRDKSNRQTIKMLVVVVLAFVLCWLPFHVGRYLQFRSLDA-PSPLLSLLSEYCSLVSVVLFYLSAAINPILYNTMSWKYRGAAARLFGLID-
Anarrhichthys ocellatus        (257) TSISSRVSHRDKSNRQTIKMLVVVVLAFVLCWLPFHVGRYVHFRSLDA-PSPLLSVLSEYCNLVSVVLFYLSAAINPILYNTMSWKYRGAAARLFGLAD-
Gasterosteus aculeatus         (257) TSISSRVSHRDKSNRQTIKMLVVVVVAFVLCWLPFHVGRYVQFRSLDN-PSPLLSVLSEYCQLVSVVLFYLSAAINPILYNTMSWKYRGAAARLFGMAD-
Pungitius pungitius            (253) TSISSRVSHRDKSNRQTIKMLVVVVLAFVLCWLPFHVGRYVHFRSLDN-PSPLLSVLSEYCNLVSVVLFYLSAAINPILYNTMSWKYRGAAARLFGMAD-
Cyclopterus lumpus             (260) TGISSRVAHRDKSNRQTIKMLVVVVLAFVLCWLPFHVGRYVHFRSLDA-PSPLLSVLSEYCNLVSVVLFYLSAAINPILYNTMSWKYRGAAARLFGLAD-
Etheostoma cragini             (259) TNISSRVAHRDKSNRQTIKMLVVVVLAFVLCWLPFHVGRYLQFRFLDA-PSPLLSLLSEYCSLVSVVLFYLSAAVNPILYNTMSWKYRGAAARLFGLAD-
Etheostoma spectabile          (259) TNISSRVAHRDKSNRQTIKMLVVVVLAFVLCWLPFHVGRYLQFRFLDA-PSPLLSLLYEYWSLVSVVLFYLSAAVNPILYNTMSWKYRGAAARLFGLAD-
Perca flavescens               (259) TSISSRVAHRDKSNRQTIKMLVVVVLAFVLCWLPFHVGRYLQFRSLDA-PSPLLSLLSEYCSLVSVVLFYLSAAINPILYNTMSWKYRGAAARLFGLAD-
Perca fluviatilis              (259) TSISSRVAHRDKSNRQTIKMLVVVVLAFVLCWLPFHVGRYLQFRSLDA-PSPLLSLLSEYCSLVSVVLFYLSAAINPILYNTMSWKYRGAAARLFGLAD-
Sander lucioperca              (259) TSISSRVAHRDKSNRQTIKMLVVVVLAFVLCWLPFHVGRYLQFRSLDA-PSPLLSLLSEYCSLVSVVLFYLSAAINPILYNTMSWKYRGAAARLFGLAD-
Cottoperca gobio               (257) TNISSRVAHRDKSNRQTIKMLVVVVLAFVLCWLPFHVGRYLQFRSLDA-PSPLLTLLSEYCSLVSVVLFYLSAAINPILYNTMSWKYRCAAARLFGMTD-
Gymnodraco acuticeps           (256) TTISSRVAHRDKSNRQTIKMLVVVVLAFVLCWLPFHVGRYLQFRSLDA-PSPLLSVVSEYCSLVSVVLFYLSAAINPILYNTMSWKYRCAAARLFGLTD-
Pseudochaenichthys georgianus  (256) TTISSRVAHRDKSNRQTIKMLVVVVLAFVLCWLPFHVGRYLQFRSLDA-PSPLLSLVSEYCSLVSVVLFYLSAAINPILYNTMSWKYRCAAARLFGLTD-
Notothenia coriiceps           (257) TTISSRVAVRDKSNRQTIKMLVVVVLAFVLCWLPFHVGRYLQFRSLDA-PSPLLSVVSEYCSLVSVVLFYLSAAINPILYNTMSWKYRCAAARLFGLTD-
Trematomus bernacchii          (257) TTISSRVAHRDKSNRQTIKMLVVVVLAFVLCWLPFHVGRYLQFRSLDA-PSPLLSVVSEYCSLVSVVLFYLSAAINPILYNTMSWKYRCAAARLFGLTD-
Epinephelus lanceolatus        (259) TNISSRVAHRDKSNRQTIKMLVVVVLAFVLCWLPFHVGRYLQFRFLDA-PSPLLSVLSEYCSLVSVVLFYLSAAINPILYNTMSWKYRGAAARLFGLTD-
Larimichthys crocea            (258) TNINSRVAHRDKSNRQTIKMLVVVVLAFVLCWLPFHVGRYLQFRFLDA-PSPLLSVLSEYCSLVSVVLFYLSAAINPILYNTMSWKYRCAAARLFGLTD-
Sebastes umbrosus              (259) TNINSRVAHRDKSNRQTIKMLVVVVLAFVLCWLPFHVGRYLQFRSLDA-PSPLLSVLSEYCSLVSVVLFYLSAAINPILYNTMSWKYRGAAARLFGLAD-
Anabas testudineus             (257) TNSSSRVAHRDKSNRQTIKMLVVVVLAFVLCWLPFHVGRYLQFRSLDA-PSPLLSILSEYCSLVSVVLFYLSAAINPILYNTMSWKYRGAAARLFGLTD-
Betta splendens                (259) TNSSSRVSHRDKSNRQTIKMLVVVVLAFVLCWLPFHVGRYLQFRSLDA-PSPLLSALSEYCSLVSVVLFYLSAAINPILYNTMSWKYRCAAARLFGLQD-
Hippoglossus hippoglossus      (259) TSISSRVAHRDKSNRQTIKMLVVVVLAFVLCWLPFHVGRYLQFRSLDA-PSPLLSLLSEYCSLVSVVLFYLSAAINPILYNTMSWKYRGAAARLFGLAD-
Hippoglossus stenolepis        (283) TSISSRVAHRDKSNRQTIKMLVVVVLAFVLCWLPFHVGRYLQFRSLDA-PSPLLSLLSEYCSLVSVVLFYLSAAINPILYNTMSWKYRGAAARLFGLAD-
Paralichthys olivaceus         (259) TSISSRVAHRDKSNRQTIKMLVVVVLAFVLCWLPFHVGRYLQFRSLDA-PSPLLSVLSEYCSLVSVVLFYLSAAINPILYNTMSWKYRGAAARLFGLAD-
Scophthalmus maximus           (260) TNSSSRVSHRDKSNRQTIKMLVVVVLAFVLCWLPFHVGRYLQFRSLDA-PSPLLSVLSEYCSLVSVVLFYLSAAINPILYNTMSWKYRGAAVRLFGLTD-
Labrus bergylta                (252) TNISSRVAHRDKSNRQTIKMLVVVVLAFVLCWLPFHVGRYLQFRSLDD-PSPLLSVLSEYCSFVSVVLFYLSAAINPILYNTMSWKYRGAAARLFGLAD-
Notolabrus celidotus           (258) TTINSRVAHRDKSNRQTIKMLVVVVLAFVLCWLPFHVGRYLQFRSLDA-PSPLLSVLSEYCSFVSVVLFYLSAAINPILYNTMSWKYRGAAARLFGLTD-
Seriola dumerili               (260) TNISSRVAHRDKSNRQTIKMLVVVVLAFVLCWLPFHVGRYLQFRSLDA-PSPLLSVLSEYCSLVSVVLFYLSAAINPILYNTMSWKYRGAAARLFGLTY-
Seriola lalandi dorsalis       (260) TNISSRVAHRDKSNRQTIKMLVVVVLAFVLCWLPFHVGRYLQFRSLDA-PSPLLSVLSEYCSLVSVVLFYLSAAINPILYNTMSWKYRGAAARLFGLTY-
Toxotes jaculatrix             (260) TNIGSRVAHRDKSNRQTIKMLVVVVLAFVLCWLPFHVGRYLQFRSLDA-PSPLLSVLSEYCSFVSVVLFYLSAAINPILYNTMSWKYRGAAARLFGLSD-
Lates calcarifer               (260) TNINSRVAHRDKSNRQTIKMLVVVVLAFVLCWLPFHVGRYLQFRSLDA-PSPLLSVLSEYCSLVSVVLFYLSAAINPILYNTMSWKYRGAAARLFGLTD-
Xiphias gladius                (260) TNISSRVAHRDKSNIQTIKMLVVVVLAFVLCWLPFHVGRYLQFRSLDA-PSPLLSALSEYCSLVSVVLFYLSAAINPILYNTMSWKYRCAAARLFGLAD-
Mastacembelus armatus          (260) TNISSRVAHRDKSNRQTIKMLVVVVLAFVLCWLPFHVGRYLQFRSLDA-PSPLLSLLSEYCSLVSVVLFYLSAAINPILYNTMSWKYRGAAARLFGLTD-
```

```
                Homo sapiens  (343) -FSQRK----LSTLKD-ESSRAWTESSINT-----------------------------
 Acanthochromis polyacanthus  (359) SHPARGR--TASTVKG-DVSNGWTESTVSF-----------------------------
          Stegastes partitus  (358) SHPARGR--TASTVKG-DGSNGWTESTVSF-----------------------------
            Monopterus albus  (358) IHATRGR--TASTTKG-DGSNGWTESTVSF-----------------------------
      Sphaeramia orbicularis  (356) TQPPRCR--TASTLKG-DSSNGWTESTVSF-----------------------------
           Anguilla anguilla  (337) -APRRT----ASAIKS-ESSPGWTESSVSM-----------------------------
    Paramormyrops kingsleyae  (338) -GPRRT----ASTIKG-ESSPCWTESGVSM-----------------------------
   Erpetoichthys calabaricus  (340) -TPKRT----ASTMKD-ESSPGWTESSIST-----------------------------
        Polypterus senegalus  (340) -TPKRT----ASTMKD-ESSPGWTESSIST-----------------------------
        Lepisosteus oculatus  (342) -APRRT----ASTMKD-ESSPGWTESSVST-----------------------------
           Carassius auratus  (337) -TPRRS----TSVVKG-ESSPCWTESTASL-----------------------------
               Danio rerio-a  (337) -IPRRS----TSVAKG-ESSPCWTESTASL-----------------------------
         Pimephales promelas  (337) -NPRRR----TSVAKG-ESPPCWTESTASV-----------------------------
               Chanos chanos  (337) -TQRRT----ASSIKG-ESSPGWTESSVSL-----------------------------
    Electrophorus electricus  (337) -VPRKH----ASAAKG-ESSPGLTESSISL-----------------------------
             Clupea harengus  (331) -IPRRT----ASSVKD-ESSPGLTESSVSL-----------------------------
        Denticeps clupeoides  (331) -APRRT----LSTAKG-ESSPGFTESSVSL-----------------------------
         Callorhinchus milii  (331) -ASGRK----PSTVKE-ESSQAWTESSIST-----------------------------
         Latimeria chalumnae  (340) -TPRRV----PSATKE-ETLPAWTESSVIT-----------------------------
       Scyliorhinus canicula  (338) -APRRT----SSTAKN-ANCSTWTETIVST-----------------------------
         Carassius auratus-b  (342) -APGQSV---QSIVNA-DSVLVWNEYSWST-----------------------------
               Danio rerio-b  (341) -APGRSL---QSIVNA-ESVSVWNEYSWST-----------------------------
         Ictalurus punctatus  (340) -TQER-----SKSLNS-ENCPVWNESSGTT-----------------------------
        Scleropages formosus  (333) -APRR----TASGVKG-DSSLGCAESSVSL-----------------------------
          Astyanax mexicanus  (362) ----RQR--RGASVTRAESCPGLNESTVSL-----------------------------
       Pygocentrus nattereri  (355) ----RRR--RSASVTKAESCPGLNESTVSLGFKEIILTAESATPRTTTLPVSLVSFHLL
 Pangasianodon hypophthalmus  (339) --------------------PSFTESSISC-----------------------------
       Tachysurus fulvidraco  (338) --------------------SSFTESSISC-----------------------------
                 Esox lucius  (365) TQPPRGR--TASTVKG-ESSPAWTESTVSL-----------------------------
         Oncorhynchus mykiss  (361) TQPPRGR--TASTVKG-ESSPAWTESTVSL-----------------------------
    Oncorhynchus tshawytscha  (361) TQPPRGR--TASTVKG-ESSPAWTESTVSL-----------------------------
                 Salmo salar  (361) TQPPRGR--TASTVKG-ESSPAWTESTVSL-----------------------------
          Salvelinus alpinus  (361) TQPPRGR--TASTVKG-ESSPAWTESTVSL-----------------------------
Boleophthalmus pectinirostris (342) NQPHRCR--TASNLKG-QCSNIWNESTASF-----------------------------
Periophthalmus magnuspinnatus (342) NQPHRCR--TASNLKG-QCSNIWNESTASF-----------------------------
                Gadus morhua  (352) GPRPRGR--TASTLKGEGCSPGWRESTVSL-----------------------------
           Takifugu rubripes  (354) SHPQRGR--TASTVKG----EGWTESTVSF-----------------------------
          Gouania willdenowi  (361) NQPPRGR--TASTLKG-DGFPGWSESTVSF-----------------------------
          Salarias fasciatus  (355) SQPARGR--TASTVKG-DGSSLWTESTVSF-----------------------------
     Austrofundulus limnaeus  (353) SQPSRCR--TASTLKG-DSSYGWTESTVSL-----------------------------
     Kryptolebias marmoratus  (376) NQPSRCR--TASTLKG-DGSYGWTESTVSL-----------------------------
         Nematolebias whitei  (353) KQPSRCR--TASSLKG-DSSCVWAESTVSL-----------------------------
         Cyprinodon tularosa  (347) GQPSRCR--TASTLKG-DSSNGWTESTVSL-----------------------------
       Cyprinodon variegatus  (347) GQPSRCR--TASTLKG-DSSNGWTESTVSL-----------------------------
       Fundulus heteroclitus  (353) GQPSRCR--TASTLKG-DSSNGWTESTVSF-----------------------------
             Poecilia formosa (358) GQPSRCR--TASTLKG-DGSNGWTESTVSF-----------------------------
            Poecilia mexicana (358) GQPSRCR--TASTLKG-DGSNGWTESTVSF-----------------------------
           Poecilia latipinna (358) GQPSRCR--TASTLKG-DGSNGWTESTVSF-----------------------------
          Poecilia reticulata (358) GQPSRCR--TASTLKG-DGSNGWTESTVSF-----------------------------
       Xiphophorus couchianus (358) GQPSRCR--TASTLKG-DGSSGWTESTVSF-----------------------------
          Xiphophorus hellerii (358) GQPSRCR--TASTLKG-DGSSGWTESTVSF-----------------------------
        Xiphophorus maculatus (358) GQPSRCR--TASTLKG-DGSSGWTESTVSF-----------------------------
       Nothobranchius furzeri (357) SQPSRCR--TASTLKG-DGSNGWTESTVSL-----------------------------
              Oryzias latipes (383) NHPARGR--TASTVKG-DSSIGWTESTISL-----------------------------
           Oryzias melastigma (384) SHPARGR--TASTVKG-DSSIGWTESTISL-----------------------------
       Cynoglossus semilaevis (363) NQPPRGRTVTASSVKG-DGLNVWTESTVSF-----------------------------
            Hippocampus comes (358) QQLPRTR--TASALKA--VADGWMESSVSF-----------------------------
              Syngnathus acus (355) QKLPRTR--TASTLKA--VTEGGMESNSF------------------------------
            Parambassis ranga (357) GQPPRGR--TASTVKA-DALNGWNESTVSF-----------------------------
     Archocentrus centrarchus (386) SLPPRGR--TASTVKG-DGSNGWTESTISF-----------------------------
     Astatotilapia calliptera (383) SLPPRGR--TASTVKG-DGSNGWTESTISF-----------------------------
          Pundamilia nyererei (383) SLPPRGR--TASTVKG-DGSNGWTESTISF-----------------------------
     Neolamprologus brichardi (358) SLPPRGR--TASTVKG-DGSNGWTESTISF-----------------------------
              Maylandia zebra (358) CLPPRGR--TASTVKG-DGSNGWTESTISF-----------------------------
        Simochromis diagramma (383) SLPPRGR--TASTVKG-DGSNGWTESTISF-----------------------------
         Haplochromis burtoni (383) SLPPRGR--TASTVKG-DGSNGWTESTISF-----------------------------
           Oreochromis aureus (383) SLPPRGR--TASTVKG-DGSNGWTESTISF-----------------------------
        Oreochromis niloticus (383) SLPPRGR--TASTVKG-DGSNGWTESTISF-----------------------------
     Thalassophryne amazonica (356) SQNPRSR--TASTVKA-DGSTGWTESTASF-----------------------------
           Echeneis naucrates (359) SQPHRGR--TASTMKG-DGSNGWTESTVSL-----------------------------
          Myripristis murdjan (353) SQPPRCR--TASTVKG-EGSPGWTESTVSL-----------------------------
          Acanthopagrus latus (359) SQPPRGR--TASTVKG-DGSNGWTESTISF-----------------------------
                Sparus aurata (358) SQPPRGR--TASTMKG-DGSNGWTESTISF-----------------------------
             Morone saxatilis (358) SQPPRGR--TASTMKG-DGSNGWTESTVSF-----------------------------
        Micropterus salmoides (358) SQPPRGR--TASTVKG-DGSNGWTESTVSF-----------------------------
      Anarrhichthys ocellatus (355) SQPPRGR--TASSMKG-DGSNGWTESTVSF-----------------------------
       Gasterosteus aculeatus (355) NQPPRGR--TASSMKG-DGSNGWTESTVSF-----------------------------
          Pungitius pungitius (351) NQQPRGR--TASSMKGGDGSNGWTESTVSF-----------------------------
           Cyclopterus lumpus (358) SQPPRGR--TASSVKG-DGSNGWTESTVSF-----------------------------
            Etheostoma cragini (357) SQPSRGR--TASTMKG-DGSNGWTESTVSF-----------------------------
        Etheostoma spectabile (357) SQPSRGR--TASTMKG-DGSNGWTESTVSF-----------------------------
             Perca flavescens (357) SQPSRGR--TASTMKG-DGSNGWTESTVSF-----------------------------
            Perca fluviatilis (357) SQPSRGR--TASTMKG-DGSNGWTESTVSF-----------------------------
            Sander lucioperca (357) SHPSRGR--TASTMKG-DGSNGWTESTVSF-----------------------------
              Cottoperca gobio (355) SQPPRGR--TASTMKG-DGLNGWTESTVSF-----------------------------
          Gymnodraco acuticeps (354) SQPPRSR--TASTLKG-DGLNGWTESTVSF-----------------------------
```

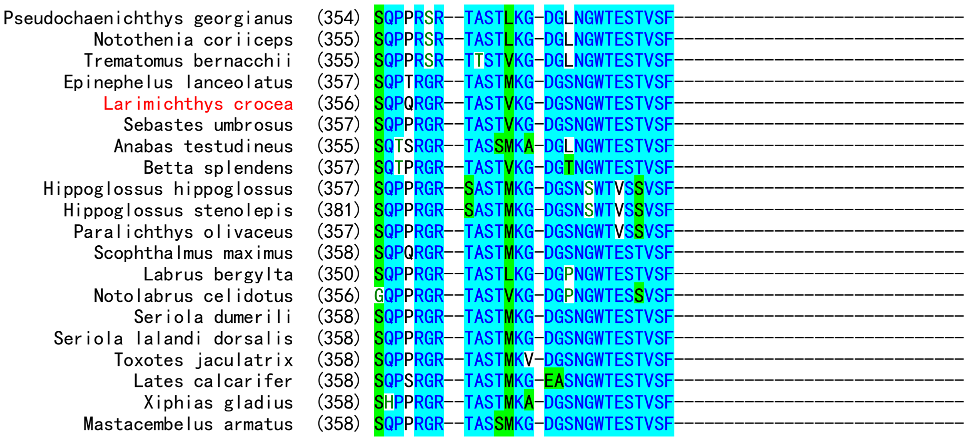


**Fig. S1.** Amino acid sequence alignment of fish GHSRs using human GHSR as a control. The species names of *Homo sapiens* (human), *Latimeria chalumnae* (coelacanth), *Danio rerio* (zebrafish), and *Larimichthys crocea* (large yellow croaker) are shown as red. The residues at 6.58 position are shown as bold letters.

### 6xHis-Dr-LEAP2

```
                              NdeI
1     ATG CAT CAC CAT CAC CAC CAT ATG GAT GAC GAT GAC AAA ATG ACC CCA CTG TGG CGT ACT GTG GGT ACG AAA CCT
      TAC GTA GTG GTA GTG GTG GTA TAC CTA CTG CTA CTG TTT TAC TGG GGT GAC ACC GCA TGA CAC CCA TGC TTT GGA
       M   H   H   H   H   H   H   M   D   D   D   D   K   M   T   P   L   W   R   T   V   G   T   K   P
                                               EK      ↑

76    CAT GGT GCG TAC TGC CAG AAC AAT TAT GAA TGT TCT ACA GGC ATT TGC CGC ATG GGC CAC TGT TCC TAC AGC CAA
      GTA CCA CGC ATG ACG GTC TTG TTA ATA CTT ACA AGA TGT CCG TAA ACG GCG TAC CCG GTG ACA AGG ATG TCG GTT
       H   G   A   Y   C   Q   N   N   Y   E   C   S   T   G   I   C   R   M   G   H   C   S   Y   S   Q

                          EcoRI
151   CCG GTT AAC TCG TAA GAA TTC
      GGC CAA TTG AGC ATT CTT AAG
       P   V   N   S   *
```

### 6xHis-La-LEAP2A

```
                              NdeI
1     ATG CAT CAC CAT CAC CAC CAT ATG GAT GAC GAT GAC AAA ATG ACC CCG CTG TGG CGT ATC ATG AAC AGT AAA CCG
      TAC GTA GTG GTA GTG GTG GTA TAC CTA CTG CTA CTG TTT TAC TGG GGC GAC ACC GCA TAG TAC TTG TCA TTT GGC
       M   H   H   H   H   H   H   M   D   D   D   D   K   M   T   P   L   W   R   I   M   N   S   K   P
                                               EK      ↑

76    TTC GGC GCT TAC TGT CAG AAT AAC TAT GAA TGC TCC ACG GGT CTG TGT CGT GCG GGT CAC TGC TCT ACT AGC CAT
      AAG CCG CGA ATG ACA GTC TTA TTG ATA CTT ACG AGG TGC CCA GAC ACA GCA CGC CCA GTG ACG AGA TGA TCG GTA
       F   G   A   Y   C   Q   N   N   Y   E   C   S   T   G   L   C   R   A   G   H   C   S   T   S   H

                                              EcoRI
151   CGC GCC ACC TCA GAG ACC GTG AAC TAC TAA GAA TTC
      GCG CGG TGG AGT CTC TGG CAC TTG ATG ATT CTT AAG
       R   A   T   S   E   T   V   N   Y   *
```

### 6xHis-La-LEAP2B

```
                              NdeI
1     ATG CAT CAC CAT CAC CAC CAT ATG GAT GAC GAT GAC AAA TCT CTG CTG TGG CGT TGG AAC AGC TTG AAG CCG GTT
      TAC GTA GTG GTA GTG GTG GTA TAC CTA CTG CTA CTG TTT AGA GAC GAC ACC GCA ACC TTG TCG AAC TTC GGC CAA
       M   H   H   H   H   H   H   M   D   D   D   D   K   S   L   L   W   R   W   N   S   L   K   P   V
                                               EK      ↑

76    GGT GCC AGT TGC CGC GAT CAC GCG GAA TGT GGC ACA AAA TAC TGC CGT AAA AAT ATT TGT TCC TTC TGG ATC AGC
      CCA CGG TCA ACG GCG CTA GTG CGC CTT ACA CCG TGT TTT ATG ACG GCA TTT TTA TAA ACA AGG AAG ACC TAG TCG
       G   A   S   C   R   D   H   A   E   C   G   T   K   Y   C   R   K   N   I   C   S   F   W   I   S

                EcoRI
151   AAC TAA GAA TTC
      TTG ATT CTT AAG
       N   *
```

### 6xHis-C4ORF48-Dr-LEAP2

```
                              NdeI
1     ATG CAT CAC CAT CAC CAC CAT ATG GAA CCG GCT GGT AGT GCC GTC CCA GCG CAG TCT CGC CCT TGC GTG GAT TGT
      TAC GTA GTG GTA GTG GTG GTA TAC CTT GGC CGA CCA TCA CGG CAG GGT CGC GTC AGA GCG GGA ACG CAC CTA ACA
       M   H   H   H   H   H   H   M   E   P   A   G   S   A   V   P   A   Q   S   R   P   C   V   D   C

76    CAT GCC TTT GAG TTC ATG CAG CGC GCA CTG CAA GAC TTG CGT AAA ACA GCC TAT AGC TTA GAT GCG CGT ACG GAA
      GTA CGG AAA CTC AAG TAC GTC GCG CGT GAC GTT CTG AAC GCA TTT TGT CGG ATA TCG AAT CTA CGC GCA TGC CTT
       H   A   F   E   F   M   Q   R   A   L   Q   D   L   R   K   T   A   Y   S   L   D   A   R   T   E

151   ACC CTT CTC CTG CAG GCA GAA CGC CGT GCC CTG TGT GCC TGC TGG CCG GCG GGC CAC GGT GGC AAA ATG ACC CCA
      TGG GAA GAG GAC GTC CGT CTT GCG GCA CGG GAC ACA CGG ACG ACC GGC CGC CCG GTG CCA CCG TTT TAC TGG GGT
       T   L   L   L   Q   A   E   R   R   A   L   C   A   C   W   P   A   G   H   G   G   K   M   T   P
                                                                                    Lys-C↑

226   CTG TGG CGT ACT GTG GGT ACG AAA CCT CAT GGT GCG TAC TGC CAG AAC AAT TAT GAA TGT TCT ACA GGC ATT TGC
      GAC ACC GCA TGA CAC CCA TGC TTT GGA GTA CCA CGC ATG ACG GTC TTG TTA ATA CTT ACA AGA TGT CCG TAA ACG
       L   W   R   T   V   G   T   K   P   H   G   A   Y   C   Q   N   N   Y   E   C   S   T   G   I   C

                                                              EcoRI
301   CGC ATG GGC CAC TGT TCC TAC AGC CAA CCG GTT AAC TCG TAA GAA TTC
      GCG TAC CCG GTG ACA AGG ATG TCG GTT GGC CAA TTG AGC ATT CTT AAG
       R   M   G   H   C   S   Y   S   Q   P   V   N   S   *
```

### 6xHis-C4ORF48-La-LEAP2A

```
                                NdeI
1     ATG CAT CAC CAT CAC CAC CAT ATG GAA CCG GCT GGT AGT GCC GTC CCA GCG CAG TCT CGC CCT TGC GTG GAT TGT
      TAC GTA GTG GTA GTG GTG GTA TAC CTT GGC CGA CCA TCA CGG CAG GGT CGC GTC AGA GCG GGA ACG CAC CTA ACA
       M   H   H   H   H   H   H   M   E   P   A   G   S   A   V   P   A   Q   S   R   P   C   V   D   C

76    CAT GCC TTT GAG TTC ATG CAG CGC GCA CTG CAA GAC TTG CGT CGC ACA GCC TAT AGC TTA GAT GCG CGT ACG GAA
      GTA CGG AAA CTC AAG TAC GTC GCG CGT GAC GTT CTG AAC GCA GCG TGT CGG ATA TCG AAT CTA CGC GCA TGC CTT
       H   A   F   E   F   M   Q   R   A   L   Q   D   L   R   R   T   A   Y   S   L   D   A   R   T   E

151   ACC CTT CTC CTG CAG GCA GAA CGC CGT GCC CTG TGT GCC TGC TGG CCG GCG GGC CAC GGT GGC AAA ATG ACC CCG
      TGG GAA GAG GAC GTC CGT CTT GCG GCA CGG GAC ACA CGG ACG ACC GGC CGC CCG GTG CCA CCG TTT TAC TGG GGC
       T   L   L   L   Q   A   E   R   R   A   L   C   A   C   W   P   A   G   H   G   G   K   M   T   P
                                                                                     Lys-C↑

226   CTG TGG CGT ATC ATG AAC AGT AAA CCG TTC GGC GCT TAC TGT CAG AAT AAC TAT GAA TGC TCC ACG GGT CTG TGT
      GAC ACC GCA TAG TAC TTG TCA TTT GGC AAG CCG CGA ATG ACA GTC TTA TTG ATA CTT ACG AGG TGC CCA GAC ACA
       L   W   R   I   M   N   S   K   P   F   G   A   Y   C   Q   N   N   Y   E   C   S   T   G   L   C

                                                                              EcoRI
301   CGT GCG GGT CAC TGC TCT ACT AGC CAT CGC GCC ACC TCA GAG ACC GTG AAC TAC TAA GAA TTC
      GCA CGC CCA GTG ACG AGA TGA TCG GTA GCG CGG TGG AGT CTC TGG CAC TTG ATG ATT CTT AAG
       R   A   G   H   C   S   T   S   H   R   A   T   S   E   T   V   N   Y   *
```

### 6xHis-C4ORF48-La-LEAP2A-srt

```
                                NdeI
1     ATG CAT CAC CAT CAC CAC CAT ATG GAA CCG GCT GGT AGT GCC GTC CCA GCG CAG TCT CGC CCT TGC GTG GAT TGT
      TAC GTA GTG GTA GTG GTG GTA TAC CTT GGC CGA CCA TCA CGG CAG GGT CGC GTC AGA GCG GGA ACG CAC CTA ACA
       M   H   H   H   H   H   H   M   E   P   A   G   S   A   V   P   A   Q   S   R   P   C   V   D   C

76    CAT GCC TTT GAG TTC ATG CAG CGC GCA CTG CAA GAC TTG CGT CGC ACA GCC TAT AGC TTA GAT GCG CGT ACG GAA
      GTA CGG AAA CTC AAG TAC GTC GCG CGT GAC GTT CTG AAC GCA GCG TGT CGG ATA TCG AAT CTA CGC GCA TGC CTT
       H   A   F   E   F   M   Q   R   A   L   Q   D   L   R   R   T   A   Y   S   L   D   A   R   T   E

151   ACC CTT CTC CTG CAG GCA GAA CGC CGT GCC CTG TGT GCC TGC TGG CCG GCG GGC CAC GGT GGC AAA ATG ACC CCG
      TGG GAA GAG GAC GTC CGT CTT GCG GCA CGG GAC ACA CGG ACG ACC GGC CGC CCG GTG CCA CCG TTT TAC TGG GGC
       T   L   L   L   Q   A   E   R   R   A   L   C   A   C   W   P   A   G   H   G   G   K   M   T   P
                                                                                     Lys-C↑

226   CTG TGG CGT ATC ATG AAC AGT AAA CCG TTC GGC GCT TAC TGT CAG AAT AAC TAT GAA TGC TCC ACG GGT CTG TGT
      GAC ACC GCA TAG TAC TTG TCA TTT GGC AAG CCG CGA ATG ACA GTC TTA TTG ATA CTT ACG AGG TGC CCA GAC ACA
       L   W   R   I   M   N   S   K   P   F   G   A   Y   C   Q   N   N   Y   E   C   S   T   G   L   C

301   CGT GCG GGT CAC TGC TCT ACT AGC CAT CGC GCC ACC TCA GAG ACC GTG AAC TAC GGT GGC GGT GGT GGC CTG CCG
      GCA CGC CCA GTG ACG AGA TGA TCG GTA GCG CGG TGG AGT CTC TGG CAC TTG ATG CCA CCG CCA CCA CCG GAC GGC
       R   A   G   H   C   S   T   S   H   R   A   T   S   E   T   V   N   Y   G   G   G   G   G   L   P

                          EcoRI
376   CGT ACC GGC GGT TAA GAA TTC
      GCA TGG CCG CCA ATT CTT AAG
       R   T   G   G   *
```

### 6xHis-Hs-LEAP2-srt

```
                             NdeI
1    ATG CAT CAC CAT CAC CAC CAT ATG GAT GAC GAT GAC AAA ATG ACC CCG TTT TGG CGT GGT GTT AGC CTG CGC CCG
     TAC GTA GTG GTA GTG GTG GTA TAC CTA CTG CTA CTG TTT TAC TGG GGC AAA ACC GCA CCA CAA TCG GAC GCG GGC
      M   H   H   H   H   H   H   M   D   D   D   D   K   M   T   P   F   W   R   G   V   S   L   R   P
                                          EK              ↑

76   ATT GGC GCG TCC TGC CGT GAT GAC TCT GAA TGT ATC ACC CGC CTG TGC CGT AAA CGC CGT TGT TCC CTG AGC GTG
     TAA CCG CGC AGG ACG GCA CTA CTG AGA CTT ACA TAG TGG GCG GAC ACG GCA TTT GCG GCA ACA AGG GAC TCG CAC
      I   G   A   S   C   R   D   D   S   E   C   I   T   R   L   C   R   K   R   R   C   S   L   S   V

                                                                 EcoRI
151  GCC CAG GAA GGA GGT GGA GGG CTG CCG CGT ACC GGT GGC TAA GAA TTC
     CGG GTC CTT CCT CCA CCT CCC GAC GGC GCA TGG CCA CCG ATT CTT AAG
      A   Q   E   G   G   G   G   L   P   R   T   G   G   *
```

**Fig. S2.** The nucleotide and amino acid sequences of LEAP2 precursors overexpressed in *E. coli*. The mature LEAP2s are shown in green. The enterokinase (EK) and Lys-C recognized residues are shaded, and their cleavage sites are indicated by arrows. The restriction enzyme cleavage sites for cloning are shaded.

### 6×His-6×Gly-NanoLuc

```
                                    NdeI
1     ATG CAT CAC CAT CAC CAC CAT ATG GAT GAC GAT GAC AAA GGT GGC GGC GGT GGT GGC AGC GGC GGT GGC GGC GGT
      TAC GTA GTG GTA GTG GTG GTA TAC CTA CTG CTA CTG TTT CCA CCG CCG CCA CCA CCG TCG CCG CCA CCG CCG CCA
       M   H   H   H   H   H   H   M   D   D   D   D   K   G   G   G   G   G   G   S   G   G   G   G   G
                                               EK        ↑

76    GGT GTC TTC ACA CTC GAA GAT TTC GTT GGG GAC TGG CGA CAG ACA GCC GGC TAC AAC CTG GAC CAA GTC CTT GAA
      CCA CAG AAG TGT GAG CTT CTA AAG CAA CCC CTG ACC GCT GTC TGT CGG CCG ATG TTG GAC CTG GTT CAG GAA CTT
       G   V   F   T   L   E   D   F   V   G   D   W   R   Q   T   A   G   Y   N   L   D   Q   V   L   E

151   CAG GGA GGT GTG TCC AGT TTG TTT CAG AAT CTC GGG GTG TCC GTA ACT CCG ATC CAA AGG ATT GTC CTG AGC GGT
      GTC CCT CCA CAC AGG TCA AAC AAA GTC TTA GAG CCC CAC AGG CAT TGA GGC TAG GTT TCC TAA CAG GAC TCG CCA
       Q   G   G   V   S   S   L   F   Q   N   L   G   V   S   V   T   P   I   Q   R   I   V   L   S   G

226   GAA AAT GGG CTG AAG ATC GAC ATC CAT GTC ATC ATC CCG TAT GAA GGT CTG AGC GGC GAC CAA ATG GGC CAG ATC
      CTT TTA CCC GAC TTC TAG CTG TAG GTA CAG TAG TAG GGC ATA CTT CCA GAC TCG CCG CTG GTT TAC CCG GTC TAG
       E   N   G   L   K   I   D   I   H   V   I   I   P   Y   E   G   L   S   G   D   Q   M   G   Q   I

301   GAA AAA ATT TTT AAG GTG GTG TAC CCT GTG GAT GAT CAT CAC TTT AAG GTG ATC CTG CAC TAT GGC ACA CTG GTA
      CTT TTT TAA AAA TTC CAC CAC ATG GGA CAC CTA CTA GTA GTG AAA TTC CAC TAG GAC GTG ATA CCG TGT GAC CAT
       E   K   I   F   K   V   V   Y   P   V   D   D   H   H   F   K   V   I   L   H   Y   G   T   L   V

376   ATC GAC GGG GTT ACG CCG AAC ATG ATC GAC TAT TTC GGA CGG CCG TAT GAA GGC ATC GCC GTG TTC GAC GGC AAA
      TAG CTG CCC CAA TGC GGC TTG TAC TAG CTG ATA AAG CCT GCC GGC ATA CTT CCG TAG CGG CAC AAG CTG CCG TTT
       I   D   G   V   T   P   N   M   I   D   Y   F   G   R   P   Y   E   G   I   A   V   F   D   G   K

451   AAG ATC ACT GTA ACA GGG ACC CTG TGG AAC GGC AAC AAA ATT ATC GAC GAG CGC CTG ATC AAC CCC GAC GGC TCC
      TTC TAG TGA CAT TGT CCC TGG GAC ACC TTG CCG TTG TTT TAA TAG CTG CTC GCG GAC TAG TTG GGG CTG CCG AGG
       K   I   T   V   T   G   T   L   W   N   G   N   K   I   I   D   E   R   L   I   N   P   D   G   S

                                                                                           EcoRI
526   CTG CTG TTC CGA GTA ACC ATC AAC GGA GTG ACC GGC TGG CGG CTG TGC GAA CGC ATT CTG GCG TAA GAA TTC
      GAC GAC AAG GCT CAT TGG TAG TTG CCT CAC TGG CCG ACC GCC GAC ACG CTT GCG TAA GAC CGC ATT CTT AAG
       L   L   F   R   V   T   I   N   G   V   T   G   W   R   L   C   E   R   I   L   A   *
```

**Fig. S3.** The nucleotide and amino acid sequence of 6×His-6×Gly-NanoLuc overexpressed in *E. coli*. The original NanoLuc is shown in blue. The enterokinase (EK) recognized residues are shaded, and it cleavage site is indicated by an arrow. The restriction enzyme cleavage sites for cloning are shaded.

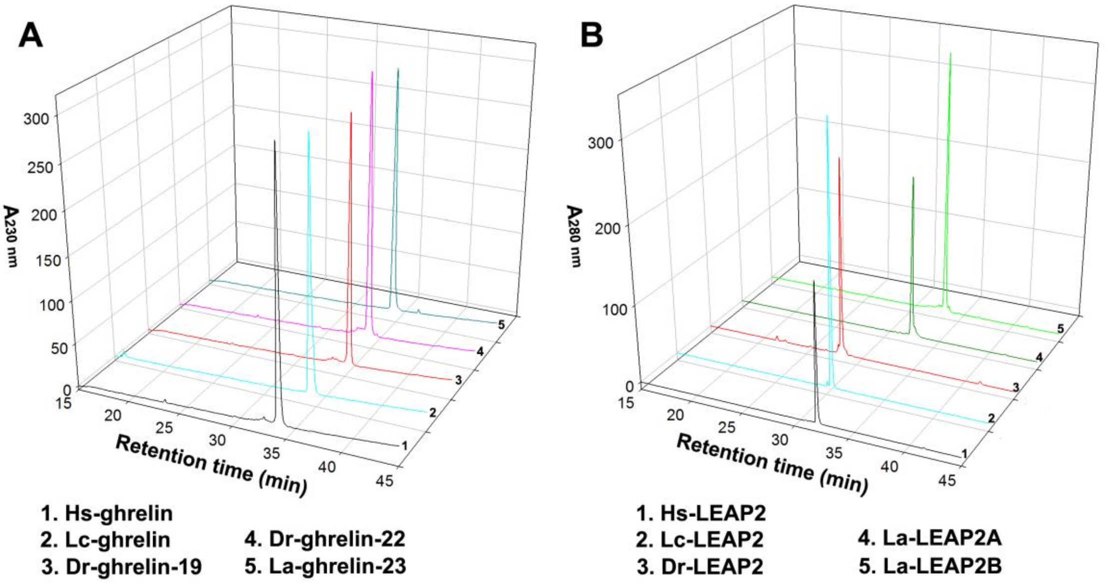


**Fig. S4.** HPLC analyses of the synthetic ghrelins (A) and recombinant LEAP2s (B). For each analysis, 15 μg of purified ghrelin or LEAP2 peptide was loaded onto an analytical C18 reverse-phase column (Zorbax 300SB-C18, 4.6 × 250 mm), and eluted from the column by an acidic acetonitrile gradient.

### Alignment of MBOAT4 (GOAT)

```
Homo sapiens          (1) --MEWLWLFFLHPISFYQGAAFPFALLFNYLCIMDSFSTRARYLFLLTGGGALAVAAMGSYAVLVFTPAVCAVALLCSLAPQQVHRWTFCFQMSWQTLCHLGLHYTEYYLH
Latimeria chalumnae   (1) ------------------------------------------YAFLLVGGFLLSNIAMGTFSLLVLIPAVLSVIMMHSISPQSVHRWVFLIQMLWQTLCHLWLQYKEHSMQ
Danio rerio           (1) MIDLLWISSDGHPQLFYQFINIPFAFLFHCLSSQGHLSIINRYVYLAMGGFMLAIATMGPYSSLLFLSAIKLLLLIHYIHPMHLHRWILGLQMCWQTCWHLYVQYQIYWLQ
Larimichthys crocea   (1) -MGLMNWLWEHHQFLMHQCFSLPFAFLFYFLAKWGYLSLRCRYLFVSIGGCVLAVVTMGIYSTLLFTSTFAFILLVCSVDSSSIHAWVFSMQMLWQTFWHLFIQYREYYLH

Homo sapiens        (110) EPPSVRFCITLSSLMLLTQRVTSLSLDICEGKVKAASGGFRSRS-SLSEHVCKALPYFSYLLFFPALLGGSLCSFQRFQARVQGSSALHPRHSFWALSWRGLQILGLECLN
Latimeria chalumnae  (70) EITRIRFIITISSLMLLTQRVTTLALDIHEGKIKIATRSSYLNN-SILQYLHNLLPYFSYMIYFPALLGGPLLSFQLFKIHIETSGVKCTENCSLPVTKKFCFFLALELLK
Danio rerio         (112) EAPDSRLLLAISALMLMTQRISSLSLDFQEG-------TISNQS--------ILIPFLTYSLYFPALLGGPLCSFNAFVQSVERQHTSMT-SYLGNLTSKISQVIVLVWIK
Larimichthys crocea (111) EPVCIRLFLAVSSLMLLTQRITSLSMDLQEKRVLTTFNALSKRNR----RV-MLLPLISYILNFTTMLGGPLCSYSRFVTLMAEIRFNHPPNPLGVVLIKLTKVMLLECLR

Homo sapiens        (220) VAVSRVVDAGAGLTDC-QQFECIYVVWTTAGLFKLTYYSHWILDDSLLHAAGFGPELGQSPGEEG--YVPDADIWTLERTHRISVFSRKWNQSTARWLRRLVFQHSRAWPL
Latimeria chalumnae (180) FLIR-NKESALTIPQS-YMINDIFVNWRTALLFKLTYYSHWILSESLLNTAGFGFKGYDNQGILVGNDLTDTDICTLETTNKISQFARAWNKTTAEWLRRLVFQRCTVHPL
Danio rerio         (207) QLFSELLKSATFNIDS--VCLDVLWIWIFSLTLRLNYYAHWKMSECVNNAAGLGVYFHKHSGQTSWDELSDGSVLVTEASSRPSVFARKWNQTTVDWLRKIVFNRTSRSPL
Larimichthys crocea (217) YCLVYFLKHNAYNPSKSIIIYGLLWVWCLALVLRIQYYSHWRISECLNNAAGFGFGENVPGVSPDWSRLSDGDFWIIEASNRMSQFARRWNATTASWLRRLVYTRCKRFPL

Homo sapiens        (328) LQTFAFSAWWHGLHPGQVFGFVCWAVMVEADYLIHSFANEFIRSWPMRLFYRTLTWAHTQLIIAYIMLAVEVRSLSSLWLLCNSYNSVFPMVYCILLLLLAKRKHKCN--
Latimeria chalumnae (289) FVTFAFSAWWHGLHPGQIFGFLFWSVAVEADKRVHYYLAPLANSWFTKQLFKVFTWIQTQLVIACIMMAIENKSFSSLWLLSQSCIILFPLLYCLVLLLLPKRPRI----
Danio rerio         (316) FMTFGFSALWHGLHPGQILGFLIWAVTVQADYKLHRFSHPKLNSLWRKRLYVCVNWAFTQLTVACVVVCVELQSLASVKLLWSSCIAVFPLLSALILIIL----------
Larimichthys crocea (328) FMTFSFSLWWHGLHLGQIVGMLTWAATVKADYHIHRCLWPKLSSSWRK-IYTCLSWINTQMIVSCIVIAVEIRNMSGLRLLSTTYIGLFPLFNIILLFILQKLKFTKEVM
```

**Fig. S5.** Amino acid sequence alignment of MBOAT4 (GOAT) from *Homo sapiens*, *Latimeria chalumnae*, *Danio rerio*, and *Larimichthys crocea*.
